# Adaptive Partitioning of the tRNA Interaction Interface by Aminoacyl-tRNA-Synthetases

**DOI:** 10.1101/312462

**Authors:** Andy Collins-Hed, David H. Ardell

**Author notes:** Email address (David H. Ardell), URL: http://davidardell.org (David H. Ardell).

## Abstract

We introduce rugged fitness landscapes called match landscapes for the coevolution of feature-based assortative interactions between *P* ≥ 2 cognate pairs of tRNAs and aminoacyl-tRNA synthetases (aaRSs) in aaRS-tRNA interaction networks. Our genotype-phenotype-fitness maps assume additive feature-matching energies, a macroscopic theory of aminoacylation kinetics including proofreading, and selection for translational accuracy in multiple, perfectly encoded site-types. We compute the stationary genotype distributions of finite panmictic, asexual populations of haploid aaRs-tRNA interaction networks evolving under mutation, genetic drift, and selection for cognate matching and non-cognate mismatching of aaRS-tRNA pairs. We compared expected genotype frequencies under different matching rules and fitness functions, both with and without linked site-specific modifiers of interaction. Under selection for translational accuracy alone, our model predicts no selection on modifiers to eliminate non-cognate interactions, so long as they are compensated by tighter cognate interactions. Only under combined selection for both translational accuracy and rate do modifiers adaptively eliminate cross-matching in non-cognate aaRS/tRNA pairs. We theorize that the encoding of macromolecular interaction networks is a genetic language that symbolically maps identifying structural and dynamic features of genes and gene-products to functions within cells. Our theory helps explain 1) the remarkable divergence in how aaRSs bind tRNAs, 2) why interaction-informative features are phylogenetically informative, 3) why the Statistical Tree of Life became more tree-like after the Darwinian Transition, and 4) an approach towards computing the probability of the random origin of an interaction network.

## 1. Introduction

Carl Woese and his co-authors argue influentially that all Earth’s cells and organelles descend not from one universal ancestor cell, but rather a communally ancestral genetic code — the one operating in ribosomal protein synthesis [1–5]. Woese’s theory is that our ancestral genetic code evolved collectively in a community of cells that exchanged genes more frequently and translated them more ambiguously than we imagine most living cells would tolerate today (although increasing the accuracy of protein synthesis can be costly, for example in bacteria competing to grow [6–9]). Our ancestral genetic code evolved as an innovation-sharing protocol [4] in a “winner-takes-all” or big bang process [10] analogous to systems competition in economics [11]. That is, the ancestral community of cells converged on one genetic code in parallel to exploit a convergently encoded pool of genes that they shared. Once enough genes came to depend on this code, and cellular fitness increasingly depended on interdependent coordination of the action of many gene products, an evolutionary phase transition occurred that “froze” the genetic code [12, 13]. In parallel, increasingly complex fitness interactions among genes, called generally *epistasis* [14, 15], cooled the rate of gene sharing, changing the evolution of cells from a genetically communal to a more vertical mode of inheritance in a Statistical Tree of Life [16], in what Woese called the *Darwinian Transition*. Broadly consistent with this theory, it was found that complexity of gene interactions (the number of pairwise interactions a gene undertakes) constrains “informational” genes from transferring horizontally between cells relative to condition-dependent “operational” genes [17, 18] and increasing pairwise protein-protein interactions, as measured in yeast two-hybrid data, reduces substitution rates in proteins [19].

In protein biosynthesis, the translation of sense codons depends directly on the identity and distribution of amino acids attached or *aminoacylated* to the 3’ ends of tRNAs at their *acceptor stems*. Aminoacylation of amino acids to tRNA acceptor stems is catalyzed in an ATP-dependent two-step reaction [20] by amino-acid-specific catalytic core domains in tRNA-binding proteins called aminoacyl-tRNA synthetases (aaRSs) [21–23]. The conserved and modular domain structure of aaRSs and the ability of some aaRSs to specifically aminoacylate model acceptor stem hairpins led to the proposal that aaRS-tRNA interactions evolved through a primordial stage of an “operational RNA code” depending on a small number of base-pairs in the acceptor stem [24, 25].

However, it is unclear how this theory fully accounts for diversity in tRNA-binding by aaRSs. As shown in Figure 1, aaRSs exhibit remarkable diversity in how they bind and interact with tRNAs. AaRSs come in two conserved and ancient superfamilies called Class I and Class II, with distinct folds, distinct mechanistic details of catalysis and — critical for our argument — distinct modes of binding to tRNAs, through opposing major or minor grooves of tRNA acceptor stems [26]. The two superfamilies may further be divided each into three subclasses [27], which pre-date the divergence of bacteria, archaea and eukarotes [23] as exemplified by the consistency with which aaRSs can be used to root the statistical Tree of Life [28]. Striking examples of aaRS pairs of different classes were found that could be docked simultaneously on tRNAs [29], which led to the hypothesis that aaRSs may have originally bound tRNAs in paired complexes to help protect tRNA acceptor stems, and subsequently diverged to single aaRS-binding with expansion of the code [30].

**Figure 1:**
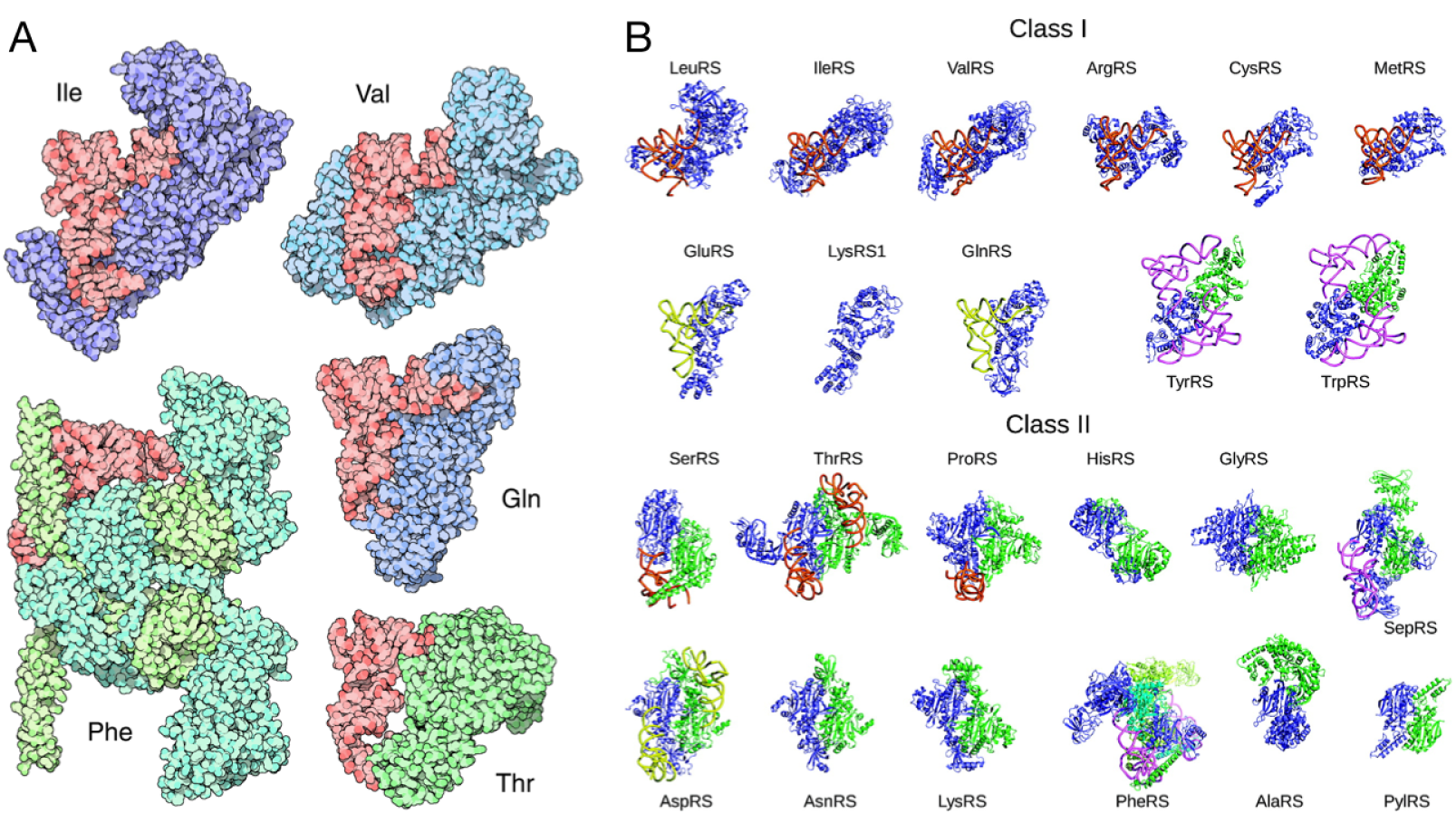
Diversity in tRNA-binding by Class I and Class II aaRSs and within aaRS subclasses. Panel A, reproduced from [31] without modification under License CC-By 4.0.: Shown in red are different species of tRNAs, all oriented identically. Aminoacyl-tRNA synthetases are shown in purple and green. Class I aaRSs, such as IleRS, ValRS and GlnRS, and class II aaRSs, such as PheRS and ThrRS, bind tRNAs on opposite faces and catalyze aminoacylation on different carbons of the last tRNA base, A76 (in Sprinzl standard coordinates [32]). Panel B: Gallery of aaRS structures co-complexed with tRNAs when available, reprinted (adapted) with permission from [33]. Copyright (2008) American Chemical Society. Subclasses a, b and c of both Class I and Class II aaRS superfamilies are indicated by orange, yellow and pink tRNA colors respectively. aaRSs are visualized with their catalytic domains in the same orientation. (two-column figure)

Because all tRNAs conform to a universal structure, tRNAs must distinguish themselves to specific aaRSs through interaction-determining features called tRNA identity elements, which vary over the major domains of life [34]. We say that the functional identity of a tRNA determines its assortative interaction with proteins as mediated by mutually compatible structural and dynamical features. Earlier, we applied an information theoretical approach to predict tRNA identity elements [35]; we call the features we predict *tRNA Class-Informative Features (CIFs*) (they could perhaps more specifically be called *Interaction-Informative Features (IIFs*)). Through comparative analysis of tRNA CIFs and also through our tRNA functional classifier [36], we have shown that tRNA CIFs are variable and phylogenetically informative within the major domains of life [37–39].

There is a widely perceived need for genetically explicit models to investigate theories about the origin and evolution of the aaRS-tRNA interaction network. For example, Vetsigian et al. [4] showed that horizontal gene transfer of protein-coding genes across a structured population of evolving codes improves the error-minimizing optimality of genetic codes, but they were unable to model the effect of horizontal transfer of components of the translational apparatus itself. They write, “a fuller account of the evolution of the genetic code requires modeling physical components of the translational apparatus, including the dynamics of tRNAs and the aminoacyl-tRNA synthetases.” Similarly, Koonin and Novozhilov [40] write, “A real understanding of the code origin and evolution is likely to be attainable only in conjunction with a credible scenario for the evolution of the coding principle itself and the translation system.” Having code evolution models with explicit evolutionary dynamics for tRNA and aaRS genes would help test other hypotheses including the roles of duplication and divergence of tRNA and aaRS genes in codon assignments [41], and even the dynamics of antisense-encoded aaRSs according to the Rodin-Ohno hypothesis [42–44].

In this work we introduce a theory for feature-based encoding of aaRS-tRNA interactions that helps answer the following questions:

1. Why are interaction-informative features phylogenetically informative?
2. How do interaction-determining features evolve and diverge while still strongly selected for function and fitness?
3. Why did more than one superfamily of aaRSs evolve with such different modes of binding tRNAs? Why is there such diversity in aaRS-binding of tRNAs even within subclasses (Fig. 1)?
4. What caused the Darwinian Transition to a more tree-like Statistical Tree of Life?
5. What is the probability of random origin of an aaRS-tRNA network of a given size?

At the outset, we considered that divergence in tRNA-binding by aaRSs could provide increased *robustness* [45] in translational accuracy to mutations in tRNAs and aaRSs (i.e. “survival of the flattest” [46]), or potentially could have evolved to increase the *evolvability* of new aaRS-tRNA interactions. Yet in the results we report here, we show that neither evolutionary robustness nor increased evolveability is necessary to positively select for divergence in tRNA-binding by aaRSs. Furthermore, selection on translational accuracy alone was insufficient to select for divergence in tRNA-binding. We found that combined selection on both accuracy and rate was necessary and sufficient for aaRS genes to evolve to adaptively partition the tRNA interaction interface. Our results depend on assumptions and modeling concepts as briefly introduced in the remainder of this section.

### 1.1. Additivity of macromolecular interaction energies

We assume that tRNAs and aaRSs interact through sets of paired features that contribute additively to their overall binding energy as manifested through dissociation rate constants. This assumption has long featured in models of DNA-protein interactions in transcription factor binding sites and their evolution [47–52] as well as on the structure and evolution of protein-protein interaction networks [53–55]. Such studies have also used the abstraction of working with simplified binary genotypes as we do here.

### 1.2. Kinetic proofreading

Hopfield [56] and Ninio [57] were the first to demonstrate the fundamental mechanism of *kinetic proofreading* now shown to underlie the accuracy of information transduction in biopolymerization reactions such as aminoacyl-tRNA selection by the ribosome [58], tRNA selection in aminoacylation [59], and nucleotide selection in transcription [60], but also cellular signal transduction [61] (but see *e.g.* [62]), including T-cell activation [63] and recently, morphogenesis [64]. In kinetic prooreading, the dissipation of cellular free energy coupled to internal, allosteric non-reactive state transitions amplifies the kinetic discrimination of substrates at some combined expense of overall reaction rate, energy, and the stochastic discard of partially processed preferred substrates [65, 66]. The discovery of proofreading was motivated in part by the observation that the amino acid selectivities of aaRSs are greater than can be explained by differences in free-energy of binding of different amino acids [56]. Ehrenberg and Blomberg [67] first derived the thermodynamic limits of kinetic proofreading in terms of the displacement from thermodynamic equilibrium of high energy cofactors such as ATP or GTP, as discussed by Kurland [68]. The kinetics of the two aaRSs classes is different; In class I aaRSs, product release is rate-limiting, while in Class II aaRSs, aminoacyl transfer is rate-limiting [69]. However, a range of different regimes of kinetic rates and allosteric state transition networks can exhibit proofreading [65, 70]. In this work we use theoretical bounds for proofreading over all possible schemes to derive bounds on aminoacylation rates.

### 1.3. Rugged landscapes, epistatic gene interactions, and modifier models

Fitness landscapes, introduced by Sewall Wright [71], map genotypes to fitnesses either directly, or via phenotypes, as recently reviewed by Ahnert [72]. Interactions between genes can cause double, triple, *etc.* mutants to have greater or lesser fitness than expected from the isolated fitness effects of their component mutations, a phenomenon known as *epistasis*. Epistasis can take place across the genotype-phenotype map at multiple scales of biological organization simultaneously [73]. Reciprocal sign epistasis (in which recombinants of haplotypes have lower fitness than non-recombinants) is a necessary (but not sufficient [74]) condition for fitness landscapes to become *rugged* [75], exhibiting potentially many separated local fitness maxima. Abstract genotype-fitness and genotype-phenotype-fitness models, such as the tunably rugged NK model [76, 77] or other regulatory or metabolic network evolution models [78–80] typically lack a concrete, mechanistic interpretation for how epistasis actually manifests through the combined actions of genes on the basis of sequences.

In this work, we allow the availability of sites for matching or mismatching between tRNA and aaRS gene products to evolve under direct genetic control, making epistasis evolveable at site resolution. As such, our work is related to population genetic models that study the genetic modification of evolutionary forces such as mutation, recombination or epistasis, called *modifier models*. Modifier models encode evolutionary parameters at neutral loci that co-evolve under uniform genetic dynamics as other *major loci* that directly impact fitness. Original applications of modifier models were aimed at studying the evolution of recombination [81, 82]. Under very general conditions near an equilibrium under viability selection, modifier loci evolve to reduce rates of mutation, migration, or recombination [83]. Without recombination, near mutation-selection balance, modifiers that increase positive or antagonistic epistasis will evolve, increasing the robustness of hap-loid asexual populations to mutations [14, 15], this robustness is an intrinsic property of the fitness landscape [84, 85]. An analysis of *fitness valley crossing* in asexual haploid populations with reciprocal sign epistasis [86] points to the critical role of the high-dimensional structure of fitness landscapes in determining evolutionary outcomes [87].

### 1.4. Origin-fixation formalism for evolutionary genetics

We model evolutionary dynamics in finite, haploid asexual populations of aaRS-tRNA networks using the *statistical mechanical* or *sequential fixation* Markov chain [49, 88], a variety of *origin-fixation model* [89] that assumes a maximum of two genotypes segregating in a population at a given time. Thus, it is assumed that the mutation rate is much smaller than the reciprocal of the square of the population size [89]. These assumptions yield an exact solution of the stationary distribution of fixed genotypes in finite populations of constant size experiencing selection, mutation and genetic drift [88, 90]. Models of this kind have been used to highlight the role of compensatory evolution on the complex genotype-phenotype-fitness landscapes of transcription-factor binding sites [50] and proteins [91]. In an appendix, Sella [90] shows results for the stationary genotype distribution of a population of haploid binary genomes selected to maximize their weight (in the coding theory sense), that is, to become “all ones.” In the Discussion, we return to this model as a natural modeling complement to the binary match landscape models that we introduce here.

## 2. Match landscapes: Models and Results

### 2.1. General Assumptions of the Current Work

Unless otherwise noted, genotypes 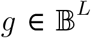 are haploid binary strings of length *L* that undergo point mutation, selection and genetic drift in panmictic Moran [93] populations of constant size — but experiencing neither recombination, duplication, deletion, insertion, inversion, conversion, nor drive (nor implicitly, horizontal transfer) of genes or genomes. We define here various *match fitness landscapes* that map genotypes 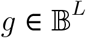 to real-valued fitnesses *w*(*g*) with 0 ≤ *w*(*g*) ≤ 1. The fitness functions we define all have at their roots a *matching function* that maps genotypes to *match matrices*, which unambiguously predict the intensities at which pairs of tRNA and aaRS species interact in a model cell or cytoplasmic volume. Except in subsection 2.7, each genotype 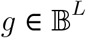 expresses an equal number *P* species of tRNA and *P* species of aaRS.

Any species of tRNA can potentially match any species of aaRS through an interaction interface shared by all. Each species of tRNA or aaRS contains the same number of sites in this shared interaction interface. A correspondence exists that partitions sites in the same way across all species, and thereby limits the way in which matches of species can occur. We call the union of single sites over all species that can potentially match or mismatch within any possible aaRS-tRNA species pair a *site-block*. Matching occurs exclusively within site-blocks, and matching is additive over site-blocks. We denote the number of site-blocks *n* and call it the *width* of the interaction interface.

### 2.2. Overview of Models and Results

A list of symbols and parameter values is given in Table 1. In subsection 2.3 we define a model we call the binary interaction channel with one site-block and compute its average fitness, load and epistasis under two different matching rules. In section 2.4, we define the P-ary interaction channel with multiple site-blocks, while in section 2.5 we present a result about its stationary genotype frequency distribution when fitness is multiplicative over site-blocks. In section 2.6 we develop an additive interaction model for aaRSs and tRNAs. In section 2.7 we re-derive a macroscopic model of aminoacylation kinetics in an interaction network with *N* tRNA species and *M* aaRS species. In section 2.8 we present results on the dependency of fitness maxima and fixed drift load on the number of cognate pairs encoded in an aaRS-tRNA network. In section 2.9 we compare fitnesses and the stationary expected frequency of masking in networks selected for translational accuracy alone versus networks selected for both accuracy and rate.

**Table 1:**
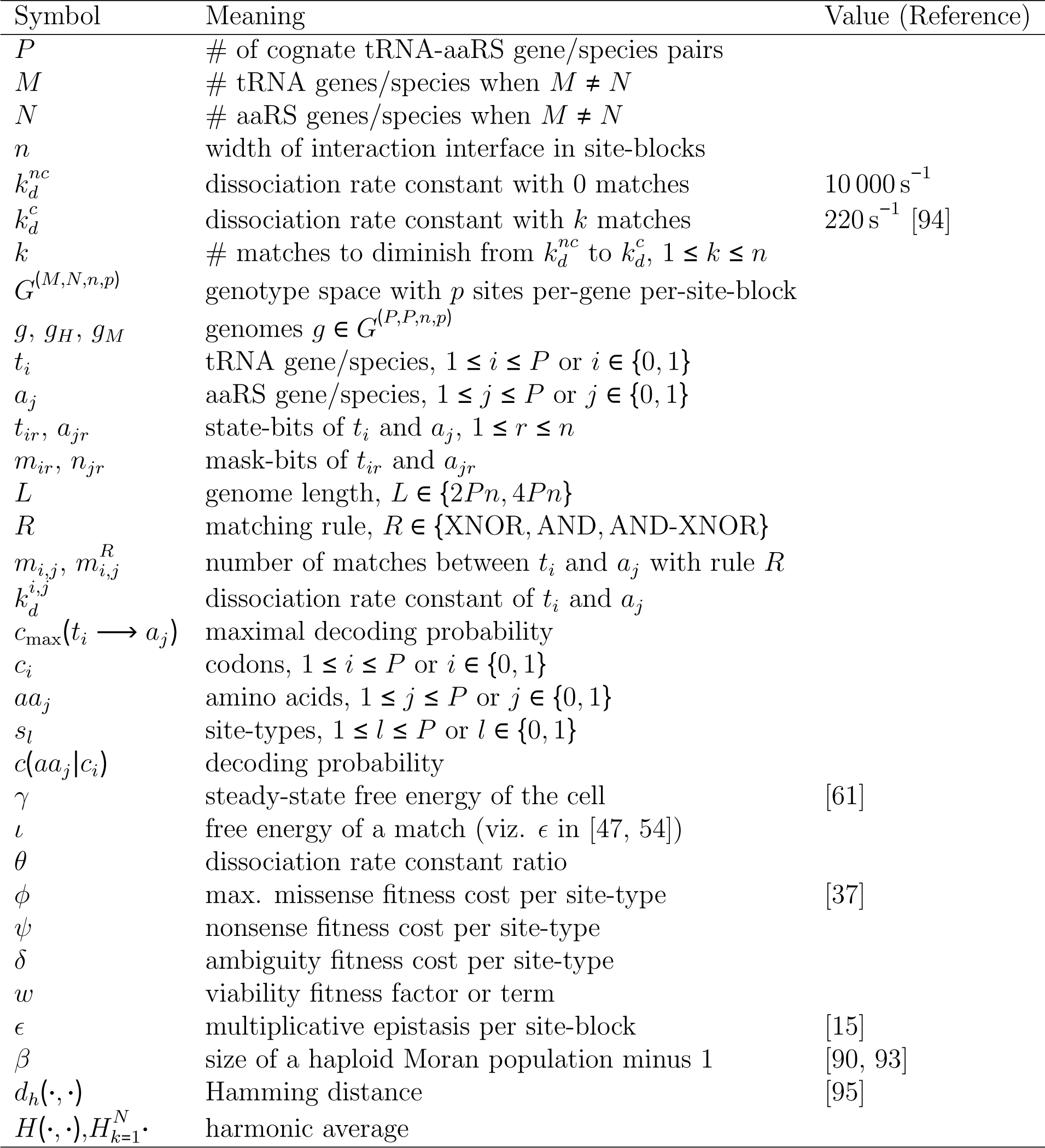
Symbols, parameter values and references for the present work.

### 2.3. The Binary Interaction Channel with an Interface of One Site-Block

Suppose that exactly one binary site in a gene for one tRNA species, *t*_0_, and another site in a gene for one aaRS species, *a*_0_, are selected to match each other, so that genotypes 11 and 00 have equal and maximal viabilities greater than those of genotypes 10 and 01, *w*_00_ = *w*_11_ > *w*_01_ = *w*_10_. This landscape is an example of “reciprocal sign epistasis.” [74, 86, 87]. In another landscape, one genotype, say 11, has higher viability than the other three, with *w*_11_ > *w*_10_ = *w*_01_ = *w*_00_. This landscape is an example of positive or antagonistic epistasis [14], in which the fitness cost of the double mutant is less than either the sum or product of the costs of single mutants. The evolution of two-locus, two-allele models has been studied under very general settings, in haploid and diploid populations with and without recombination and modifiers of epistasis, most recently in the haploid setting by Liberman and Feldman [15]. The minimal setting for a binary feed-forward interaction channel, encoding up to two amino acids, is only slightly more complex than the two-locus, two-allele model. It is a four-locus, two-allele model representing genes for two tRNA species *t*_0_ and *t*_1_ and two aaRS species *a*_0_ and *a*_1_, in which either tRNA can potentially match either aaRS through a single site-block. Depending on the matching rule and the specific genotype, either of the two tRNA species may match zero, one or both aaRS species.

We define two different matching rules in our model through logical operations on bits. The first we call the *XNOR rule* and indicate it in Table 2 and elsewhere with the ⟺ symbol. Using the XNOR rule, the match score 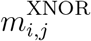 of *t*_*i*_ and *a*_*j*_, with *i,j* ∈ {0,1} is:

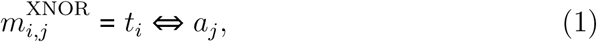

where (*a* ⟺ *b*) ≡ (*a* ⊙ *b*) ≡ ¬(*a* ⊕ *b*) is the logical XNOR of *a* and *b*.

**Table 2:**
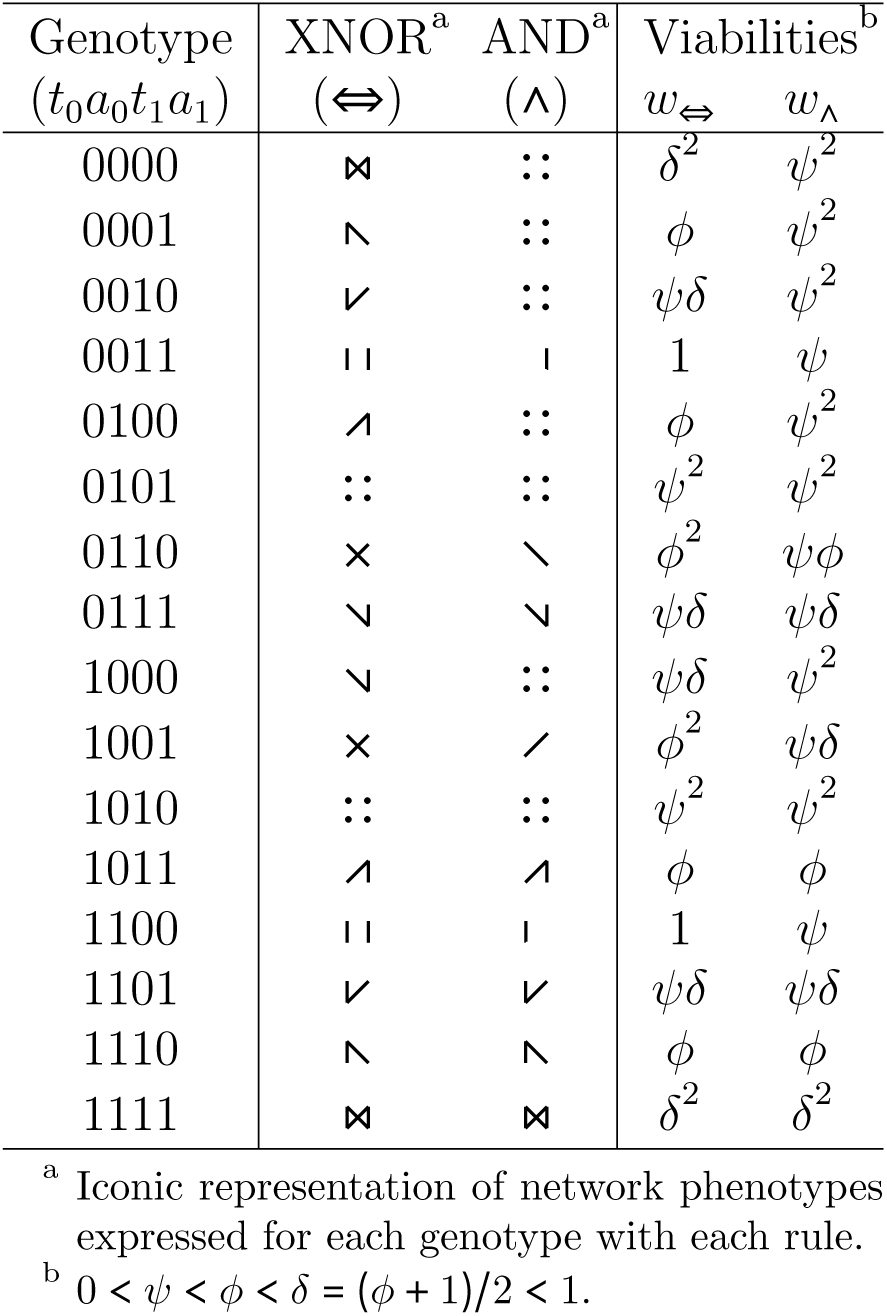
Viabilities of the symmetric multiplicative binary interaction channel with one site-block under two different matching rules.

The second we call the *AND rule* and indicate it in Table 2 and elsewhere with the Λ symbol. Using the AND rule, the match score 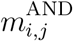 of *t*_*i*_ and *a*_*j*_, with *i*, *j* ∈ 0, 1 is:

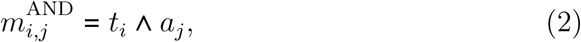

where (*a* Λ *b*) is the logical AND of *a* and *b*.

According to the set-up in Panel A of Fig. 2, we suppose that all sources of ambiguity are collected into the network. The interaction of these four species of gene products occurs through a single site for each of them. Both aaRS species have equal concentration and efficiency, both tRNA species have equal concentration and both amino acids have equal concentration. There are two equally frequent site-types *s*_0_ and *s*_1_ using the terminology and assumptions of [37, 92], one at coordinate *x*_0_ = 0 and the other at coordinate *x*_l_ = 1. Amino acids *aa*_0_ and *aa*_l_ obtain maximal viability 1 in their respective site-types *s*_0_ and *s*_1_. Amino acid *aa*_0_ obtains viability *ϕ* in site-type *s*_1_ and *vice versa*, while the viability of an unencoded amino acid (corresponding to when an aaRS species has no tRNA species that matches it) is *ψ*, with 0 < *ψ* < *ϕ* < 1. Only codons of type *c*_0_, which are exclusively and perfectly read by tRNA species *t*_0_, exist in sites of type *s*_0_, while only codons of type *c*_1_, which are exclusively and perfectly read by tRNA species *t*_1_, exist in sites of type *s*_1_. Amino acid *aa*_0_ is charged exclusively and perfectly by aaRS *a*_0_ and amino acid *aa*_1_ is charged exclusively and perfectly by aaRS *a*_1_. If a tRNA matches both aaRSs, the codons it reads achieve a fitness *δ* = (*ϕ* + 1)/2, which is the arithmetic average of its translations. Thus, ambiguity is more fit than pure missense, *δ* > *ϕ*. The fitness of a genotype is the product of its fitness in the two site-types. With these assumptions, we write the fitnesses of the 16 possible genotypes under two different matching rules in Table 2.

**Figure 2:**
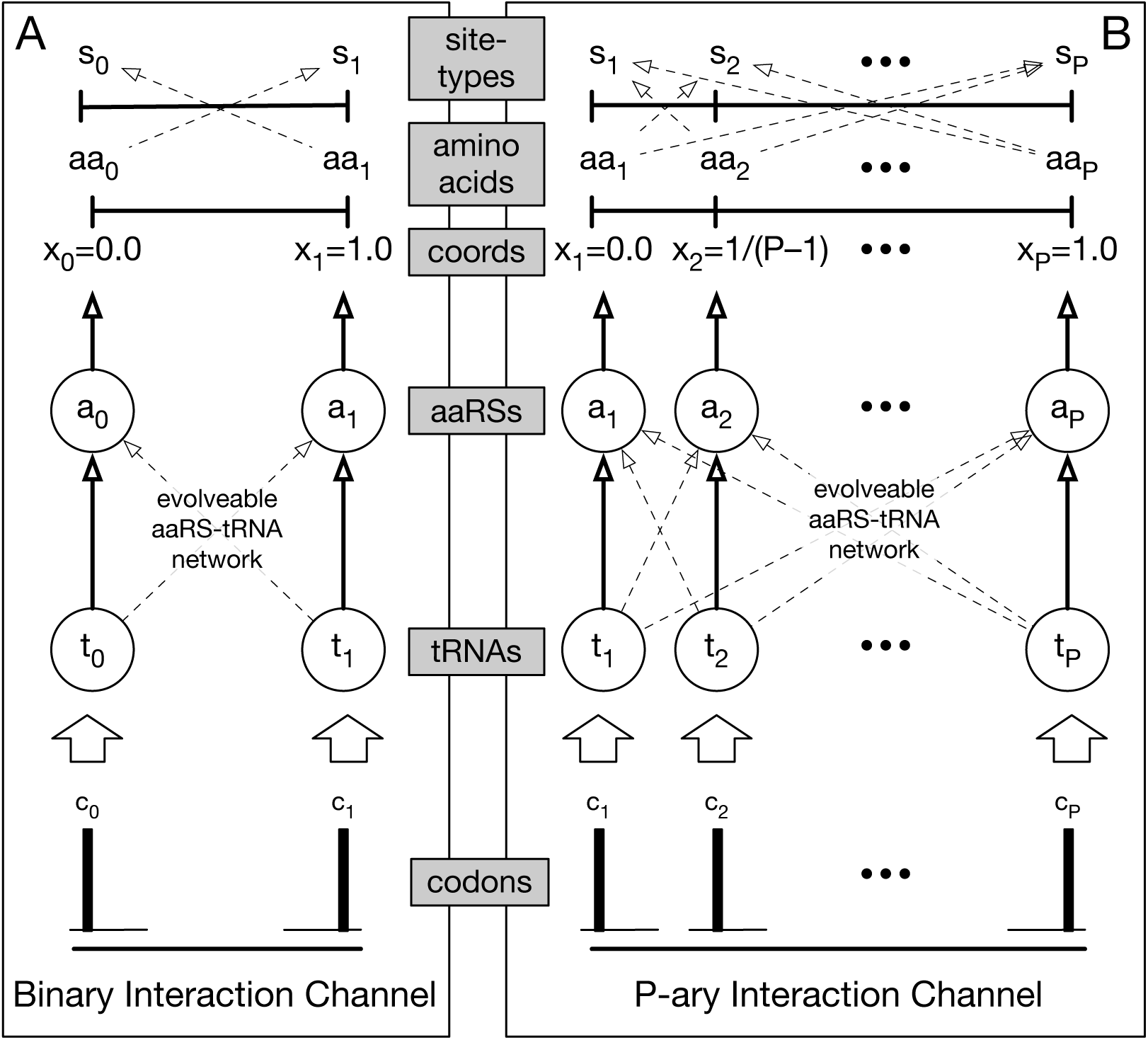
Set-up for models comparing fitness landscapes with different aaRS-tRNA networks and network encodings. Except in section 2.7, there are always a fixed and equal number *P* ≥ 2 species of tRNA, *P* species of aaRS, *P* codons, *P* available amino acids, and *P* site-types, the latter two of which are uniformly and maximally distributed within a one-dimensional amino-acid/site-type space representing differential selection on amino acid side chain properties such as hydrophobicity [92](labelled as “coords”). To each site-type corresponds a unique codon that encodes it perfectly and a unique amino acid that fits it perfectly. To each codon corresponds a unique tRNA that reads it perfectly. To each amino acid corresponds a unique aaRS that charges it perfectly. Panel A. The Binary Interaction Network Channel (*P* = 2) studied in subsection 2.3. Panel B. The P-ary Interaction Network Channel studied in subsections 2.8 and 2.9. (one-column figure)

Table 2 gives all genotype viabilities for the binary interaction channel with one site-block under the two different matching rules, XNOR and AND. The channel achieves greater maximum fitness using the XNOR rule because it can encode two interactions simultaneously with it, but only one with the AND rule. Inspecting the fitnesses of genotypes in consideration of the assumed inequality 0 < *ψ* < *ϕ* < *δ* = (*ϕ* + 1)/2 < 1, one finds that the fitness of every genotype with the XNOR rule is greater than or equal to its fitness with the AND rule. From eq. 9 in [90], one may infer directly that with these fitnesses under the stationary genotype distribution of the “sequential fixations” origin-fixation process [88, 90], the binary interaction channel has both a higher average fitness and a smaller fixed-drift load with the XNOR rule than it does with the AND rule, for all values of population size parameter *β* and for all 0 < *ψ* < *ϕ* < 1.

Liberman and Feldman [15] define multiplicative epistasis for the two-locus, two-allele model analogously to:

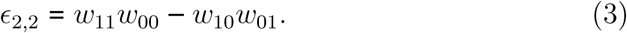

A generalization of this expression to four loci and two alleles is:

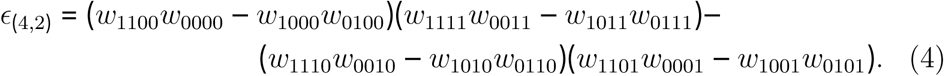

After substituting fitnesses from Table 2 and simplification, we find that the multiplicative epistasis 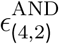 of the AND rule is always positive:

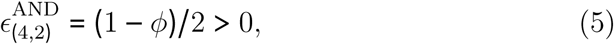

and that the multiplicative epistasis 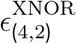 of the XNOR rule is also always positive:

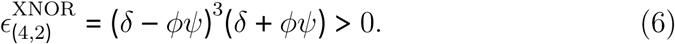

### 2.4. The P-ary Interaction Channel over an Interface of Multiple Site-Blocks

We now extend the model of section 2.3 by assuming that the interaction intensities of *P* > 2 tRNA species, labeled *t*_*i*_ with 1 ≤ *i* ≤ *P*, and *P* aaRS species, labeled *a*_*j*_ with 1 ≤ *j* ≤ *P*, depend directly on their *match scores* 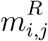 with matching rule *R*, which are additive over an interaction interface of width *n* > 1 site-blocks. To do so, we introduce two different combinations of genotype spaces and matching rules to be used in the sequel. We first define a genotype space *G*^(*P,P,n*,1)^ of dimension 2*Pn* and explain how we apply an XNOR matching function 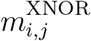 to genotypes from that space to obtain the results of section 2.8. We then define a second larger genotype space *G*^(*P,P,n*,2)^ of dimension 4*Pn* and explain how we apply a more complex matching function 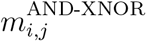 on genotypes from that space to obtain the results of section 2.9.

Assuming every species of tRNA or aaRS is produced by only one gene, we assign *n state-bits* to each of the 2*P* tRNA and aaRS genes and write them as follows: *t*_*i*_ ≡ *t*_*i*1_*t*_*i*2_ … *t*_*ir*_ … *t*_*in*_, and *a*_*j*_ ≡ *a*_*j*1_ … *a*_*jr*_ … *a*_*jn*_ respectively, where multiplication in this case implies string concatenation, 1 ≤ *i*, *j* ≤ *P*, 1 ≤ *r* ≤ *n*, and 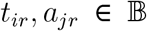. We then order and concatenate genes into genotypes as follows: *g* ≡ *t*_1_*a*_1_*t*_2_*a*_2_ … *t*_*P*_*a*_*P*_. Denote by *G*^(*P,P,n*,1)^ the set of all possible binary genotypes with *P* tRNA genes and *P* aaRS genes of width *n* site-blocks and one site per-gene per-site-block, of total length *L* = 2*Pn*. For any genotype *g* ∈ *G*^(*P,P,n*,1)^ the match score 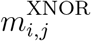 of *t*_*i*_ and *a*_*j*_ in the XNOR matching function is defined as:

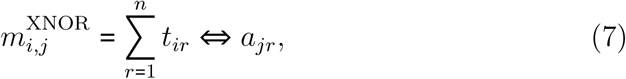

where (*a* ⟺ *b*) ≡ (*a* ⊙ *b*) ≡ ¬(*a* ⊕ *b*) is the XNOR of *a* and *b*, true when (*a* ⊕ *b*), the XOR of *a* and *b*, is false. The XNOR match score 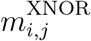 of *t*_*i*_ and *a*_*j*_ is inversely related to their Hamming distance *d*_*H*_(*t*_*i*_, *a*_*j*_):

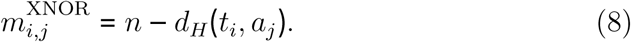

We now introduce a third matching rule, which we call the *AND-XNOR*, *MASKED-XNOR*, or *MASKED-MATCH* rule. Suppose that every macro-molecular species adds one evolveable *mask bit* that switches on or off the accessibility for matching of exactly one of its *state bits* (Fig. 3). Mask-bits are site-specific interaction modifiers. Now, with 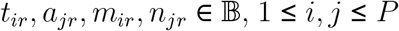 and 1 ≤ *r* ≤ *n*, we assign *n* state-bits to each of the *P* tRNA genes as before, writing the state-bits of tRNA gene *t*_*i*_ as *t*_*i*1_ … *t*_*ir*_ … *t*_*in*_, and in addition, we assign *n* mask-bits to each of the *P* tRNA genes, writing the mask-bits of tRNA gene *t*_*i*_ as *m*_*i*1_ … *m*_*ir*_ … *m*_*in*_, so that *m*_*ir*_ is the mask-bit corresponding to state-bit *t*_*ir*_. Similarly, we assign *n* state-bits to aaRS gene and write them as *a*_*j*1_ … *a*_*jr*_ … *a*_*jn*_. In addition, we assign *n* mask-bits to each of the *P* aaRS genes, and write the mask-bits of aaRS gene *a*_*j*_ as *n*_*j*1_ … *n*_*jr*_ … *n*_*jn*_, so that *n*_*jr*_ is the mask-bit corresponding to state-bit *a*_*jr*_. Finally, we order and concatenate genes into genotypes as follows (without loss of generality): *g* ≡ *t*_1_*a*_1_*t*_2_*a*_2_ … *t*_*p*_*a*_*P*_*m*_1_*n*_1_*m*_2_*n*_2_ … *m*_*P*_*n*_*P*_. Denote by *G*^(*P,P,n*,2)^ the set of all possible binary genotypes with *P* tRNA genes and *P* aaRS genes interacting over width *n* site-blocks, with 2 sites per-gene per-site-block, and a total length *L* = 4*Pn*. For any genotype *g* ∈ *G*^(*P,P,n*,2)^ the match score 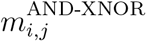 of *t*_*i*_ and *a*_*j*_ with AND-XNOR matching rule is defined:

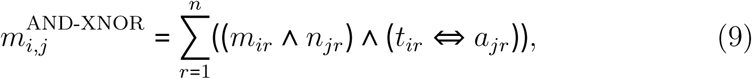

where (*a* Λ *b*) is the logical AND of *a* and *b*.

**Figure 3:**
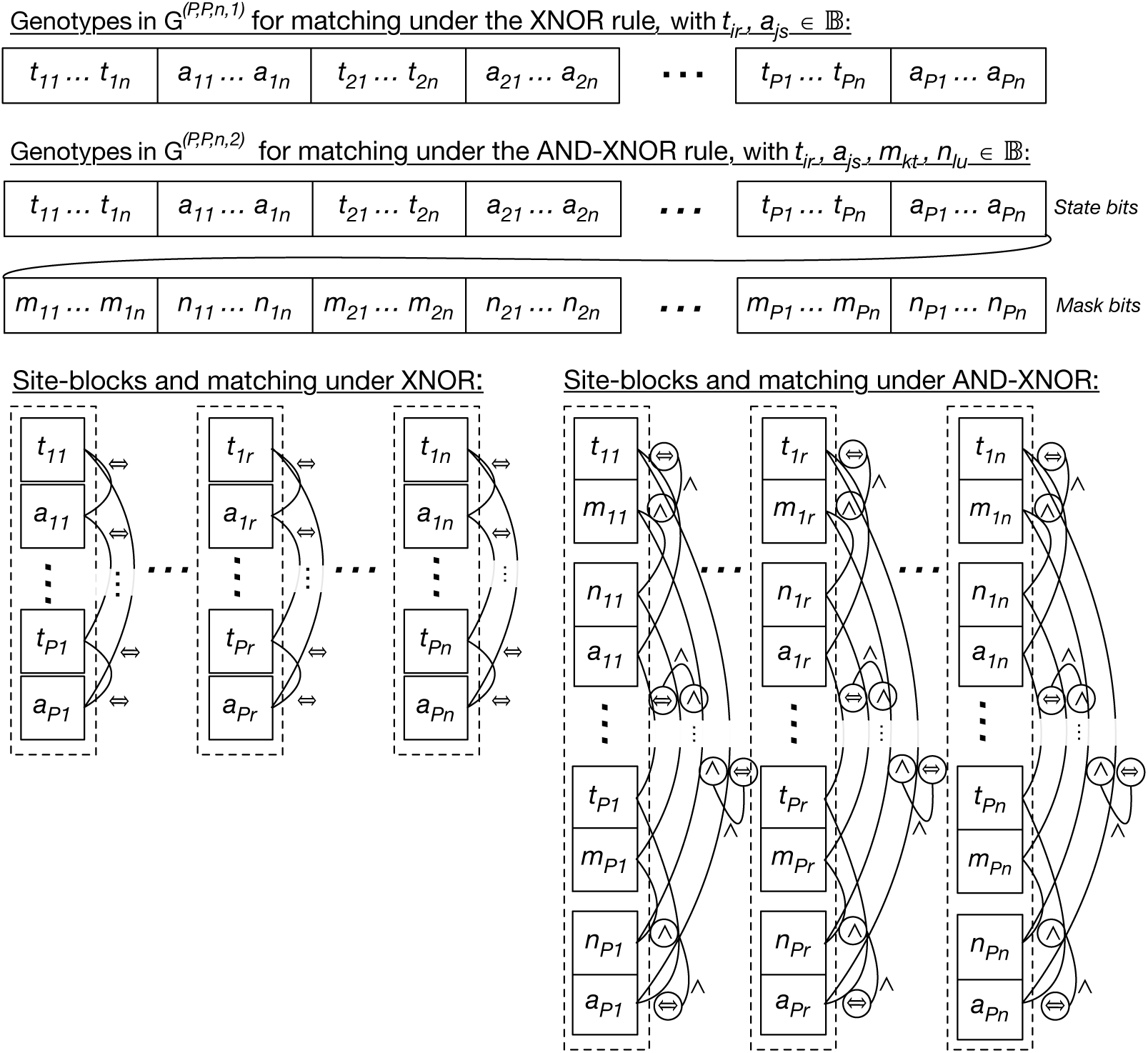
Genotype spaces, site-blocks and matching with the XNOR and AND-XNOR matching rules. (one or two-column figure)

### 2.5. P-ary interaction channels with multiplicative fitness over site-blocks

Let fitness depend multiplicatively on the match scores of corresponding tRNA, aaRS species pairs (*i.e.* those that share the same index), and inversely on the match scores of non-corresponding tRNA, aaRS species pairs (*i.e.* those with different indices). For example, if the fitness contributions of a match between any cognate pair or of mismatch between any non-cognate pair, one might define the viability fitness *w*(*g*) of genotype *g* ∈ *G*^(*P,P,n*,1)^ as:

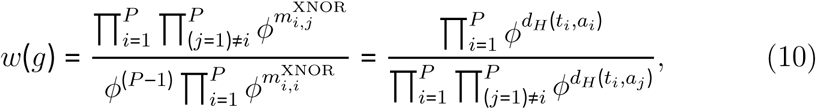

where 0 < *ϕ* ≤ 1 is a selection intensity parameter. The viabilities of eq. 10 are positive and less than or equal to 1, and increase both as tRNAs and aaRSs of the same index match while tRNAs and aaRSs of different indices mismatch. In the appendix, we show that the function in eq. 10 is multiplicative over site-blocks as previously defined, and that for all fitness functions multiplicative over site-blocks, the stationary distribution of fixed genotypes of [90] may readily be obtained as a product of the stationary frequencies of site-blocks. This result should be compared to Result 2 in [15], which states that in a large two-locus, two-allele haploid population in mutation-selection balance, a unique polymorphic equilibrium with full linkage equilibrium exists only in the absence of multiplicative epistasis.

### 2.6. From additive interaction energies to kinetic rate constants

As simple and tractable as the fitness function in eq. 10 may be, it is more realistic to suppose that the fitness of an aaRS-tRNA network is manifested through its translation of protein-coding genes. We therefore wish to create a decoding function that takes a match matrix as input and outputs a *decoding matrix* that specifies the conditional aminoacylation profile of every tRNA species.

We assume through the sequel that matches *m*_*i,j*_ between tRNA species *t*_*i*_ and aaRS species *a*_*j*_ contribute additively to their binding energy in an aaRS-tRNA complex (whether activated or not), and that only one kinetic rate constant depends on this energy and varies from complex to complex with all other kinetic rate constants set equal (see next section). Table 5 in Schimmel and Söll [94] displays kinetic data for aaRS-tRNA complexes with data from [96, 97] of about 220 s^−1^ for cognate aaRS-tRNA complexes and about 1600 s^−1^ for near-cognate interactions. We assumed a cognate dissociation rate constant of 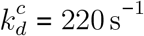 and a non-cognate dissociation rate constant of 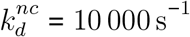 representing the background energy of interaction between tRNAs and aaRSs, also comparable to data in [98].

Define *k* as the number of matches required to diminish dissociation rate from 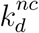 to 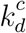, with 1 ≤ *k* ≤ n. Following Johnson and Hummer [54], we calculate non-cognate and cognate equilibrium constants as reciprocals of the non-cognate and cognate dissociation rates. The dissociation rate constant 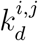 between tRNA *t*_*i*_ and aaRS *a*_*j*_ with *m*_*i,j*_ matches, 0 ≤ *m*_*i,j*_ ≤ *n* then may be defined

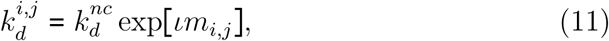

where 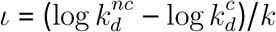.

### 2.7. Decoding functions for macroscopic, well-mixed proofreading aaRS-tRNA networks

We now assume that matching feature-set-pairs contribute additively to interaction energies between species pairs and transform interaction energies into kinetic rates of dissociation, or off-rates, of aaRS-tRNA species-pair complexes (in this section, 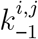 is the same as 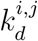 in eq. 11). We elaborate on the reaction scheme shown in fig. 4A to compute the decoding rate/aminoacylation probability of one species of tRNA *t*_0_ interacting in well-mixed volume with two species of aaRSs *a*_0_ or *a*_1_ at equal concentration as in fig. 4B. We then generalize this to calculate the *maximal decoding probability c*_max_(*t*_*i*_ → *a*_*j*_) that a tRNA of species *t*_*i*_, with 1 ≤ *i* ≤ *M* was last aminoacy-lated by an aaRS of species *a*_*j*_, with 1 ≤ *j* ≤ *N* in an aaRS-tRNA network of *M* tRNA species and *N* aaRS species with variable dissociation rate constants 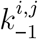 that vary between complexes of different aaRS-tRNA species pairs (fig. 4C). A comparable development was presented in [99], who were particularly interested in the energy costs of proofreading.

**Figure 4:**
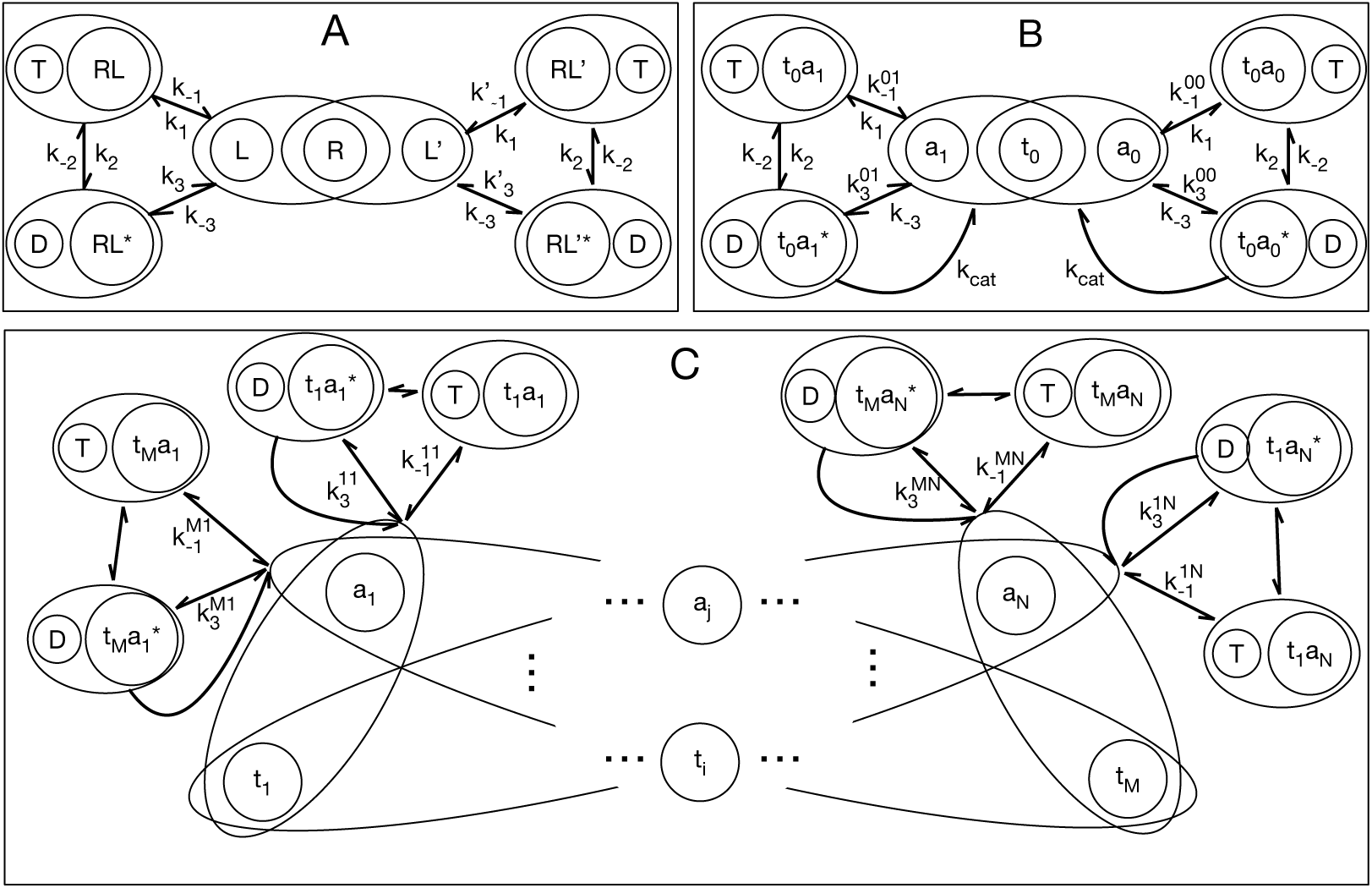
Application of kinetic proofreading schemes to compute decoding rates in a macroscopic interaction network of *M* ≥ 2 species of tRNAs and *N* ≥ 2 species of aaRSs, including kinetic proofreading of tRNAs by aaRSs but presently ignoring errors in amino acid selection by aaRSs or tRNA selection on ribosomes. A. The single-molecule two-cycle, two-state kinetic proofreading scheme of [61, fig. 1] for a receptor R that can preferentially select ligands of type *L*’ over ligands of type *L*, assuming 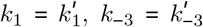 and 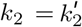 are all pseudo-first-order rate constants and the concentrations of ligand species are equal, *i.e.* [*L*] = [*L*’] ≫ [*R*]. B. The scheme from panel A redrawn from the perspective of a single tRNA species *t*_0_ alternatively aminoacylated (and instantaneously deacylated) by two aaRS species *a*_0_ and *a*_1_ of equal concentrations through catalytic steps with rate *k*_*cat*_ ≪ *k*_3_, thus [*t*_0_] ≫ [*a*_0_] = [*a*_1_] ≫ 1. C. Generalization of the scheme in B to *M* species of tRNAs and *N* species of aaRSs. All corresponding rate constants are assumed equal across all interactions except those indicated. (one-or two-column figure)

Qian [61] re-cast the classic Hopfield kinetic proofreading model as the five-state Markov Chain shown in fig. 4A, describing a cell signalling receptor *R* with a two-step activation scheme that discriminates against ligand *L* in favor of ligand *L*’ via off-rates (dissociation rates). The error rate per-receptor *ƒ* is the ratio of activated receptor affinities with ligands *L* and *L*’. Qian [61] computed the minimum error rate per-receptor *ƒ*_min_ for any set of kinetic constants in terms of the dissociation-rate-constant ratio 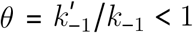 and an exponential function of the steady-state free energy of the cell γ = *e*^(Δ*G*_*DT*_/*RT*)^ ≥ 1, associated with the (deliberately unbalanced) coupled reactions *T* ⇌ *D* in fig. 4, namely 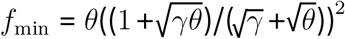. Qian [61] also re-derived the absolute lower thermodynamic limit over all possible kinetic-proofreading schemes [67], and the classical minimum per-receptor error-rate *ƒ*_min_ in the two-state kinetic proofreading scheme shown in fig. 4A, with *θ*^2^ ≤ *ƒ*_min_ ≤ *θ* [56, 57]. These two bounds correspond to perfect proofreading (with infinite ATP) on the left and thermodynamic equilibrium/recent death on the right.

These results apply equally well to enzymes as the rate of catalysis (*k*_*cat*_ in figs. 4B and 4C) vanishes. This is one of three conditions on the kinetic rate constants that achieve the minimum error rate *ƒ*_min_ [56, 61]. To achieve accuracy, enzymes and receptors add states from which they discard cognate substrates at appreciable rates so they can give non-cognate substrates more time to dissociate.

If the concentrations of aaRSs are large and equal to each other, the treatment of Qian [61] applies to Fig. 4B even though the roles of ligand and receptor are reversed. Let us define 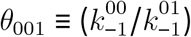 as the ratio of dissociation rate constants of tRNA *t*_0_ with aaRS *a*_0_ and aaRS *a*_1_, and similarly 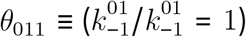. Then, at steady state, the relative rate of aminoacylation of tRNA *t*_0_ by aaRS *a*_1_ versus aaRS *a*_0_ may be written 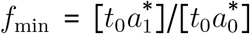, bounded by 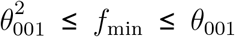 and the time-averaged maximal decoding probability *c*_max_(*t*_0_ → *a*_1_) that tRNA *t*_0_ was last aminoacylated by aaRS *a*_1_ is:

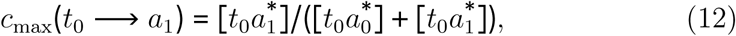

with

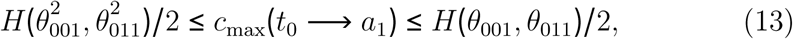

where H(*α,β*) is the harmonic average of *α* and *β*. The maximal decoding probability is maximal over all kinetic schemes of aminoacylation; however, by the data processing inequality, it is also the maximal accuracy of translation over all error-rates in tRNA-selection by ribosomes.

More generally, let us define *θ*_*ikj*_ as the ratio of dissociation rate constants of tRNA *t*_*i*_ with aaRS *a*_*k*_ and aaRS *a*_*j*_ respectively, *i.e.* 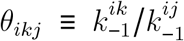, with 1 ≤ *i* ≤ *M* and 1 ≤ *j*, *k* ≤ *N*. The maximal decoding probability *c*_max_(*t*_*i*_ → *a*_*j*_), that a tRNA of species *t*_*i*_ was last aminoacylated by an aaRS of species *a*_*j*_ in an aaRS-tRNA network of *M* species of tRNA and *N* species of aaRS, is

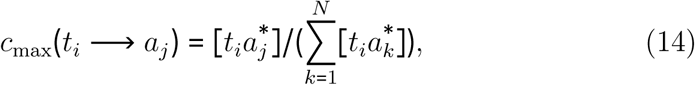

with

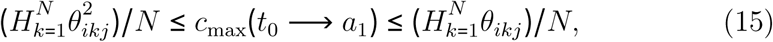

where 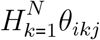 is the harmonic average over all *θ*_*ikj*_, 1 ≤ *k* ≤ *N*.

### 2.8. The Dependence of Load on Number of Encoded Amino Acids

Drawing on the terminology and concepts of earlier work [37, 92, 100, 101], we present a highly simplified translational system to compare fitnesses and stationary genotype frequencies of different matching rules. With reference to Fig. 2B, we continue to assume *P* pairs of aaRS and tRNA species, as well as *P* species of codons, amino acids, and site-types, so that tRNA species *t*_*i*_, 1 ≤ *i* ≤ *P* always reads codon *c*_*i*_, while aaRS *a*_*i*_ always charges amino acid *aa*_*i*_, which has maximal fitness in sites of type *s*_*i*_. With these assumptions, the decoding probability *c*(*aa*_*j*_ |*c*_*i*_) of decoding codon *c*_*i*_ as amino acid *aa*_*j*_ is equal to *c*_max_(*t*_*i*_ → *a*_*j*_) of the last section, *c*(*aa*_*j*_|*c*_*i*_) ≡ *c*_max_(*t*_*i*_ → *a*_*j*_). The fitness *w*(*aa*_*j*_|*s*_*l*_) of amino acid *aa*_*j*_ in site-type *s*_*l*_, with 1 ≤ *j*, *l* ≤ *P* is *ϕ*^|*x*_*j*_ − *x*_*l*_|^, where *x*_*i*_ = (*i* − 1)/(*P* − 1). The fitness *w*_*l*_ in site-type *s*_*l*_ is the expected fitness of translations of codons occupying that site-type, here exclusively codon *c*_*l*_, *i.e.* 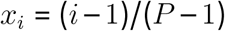. The fitness *w*_*A*_(*g*) of genotype *g* selected for translational accuracy alone is the product of its fitnesses over all site-types:

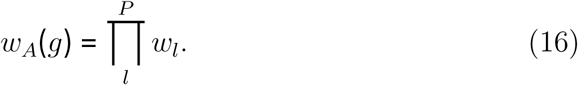

We implemented this model in a Python 3 script called “atINFLAT” for “aaRS-tRNA Interaction Network Fitness Landscape Topographer,” available as supplementary data. It can compute the stationary genotype distributions of small networks and compute statistics such as fitnesses for individual genotypes from much larger networks.

It is easy to prove that binary codes with zero matches between any codewords have a maximum size of only two codewords [102, 103]. Thus, with the XNOR rule, in which tRNAs and aaRSs may potentially match or mismatch over their entire shared interface, the interactions of only two aaRS-tRNA pairs may be encoded perfectly without cross-matching. As predicted, when we used atINFLAT to compute maximum and average fitnesses on landscapes with and without proofreading, we found that both the maximum fitness decreased and fixed-drift load increased when more than two cognate pairs were overloaded on the same interaction interface, reflecting an increasing cost of translational missense as more amino acids get encoded (Fig. 5).

**Figure 5:**
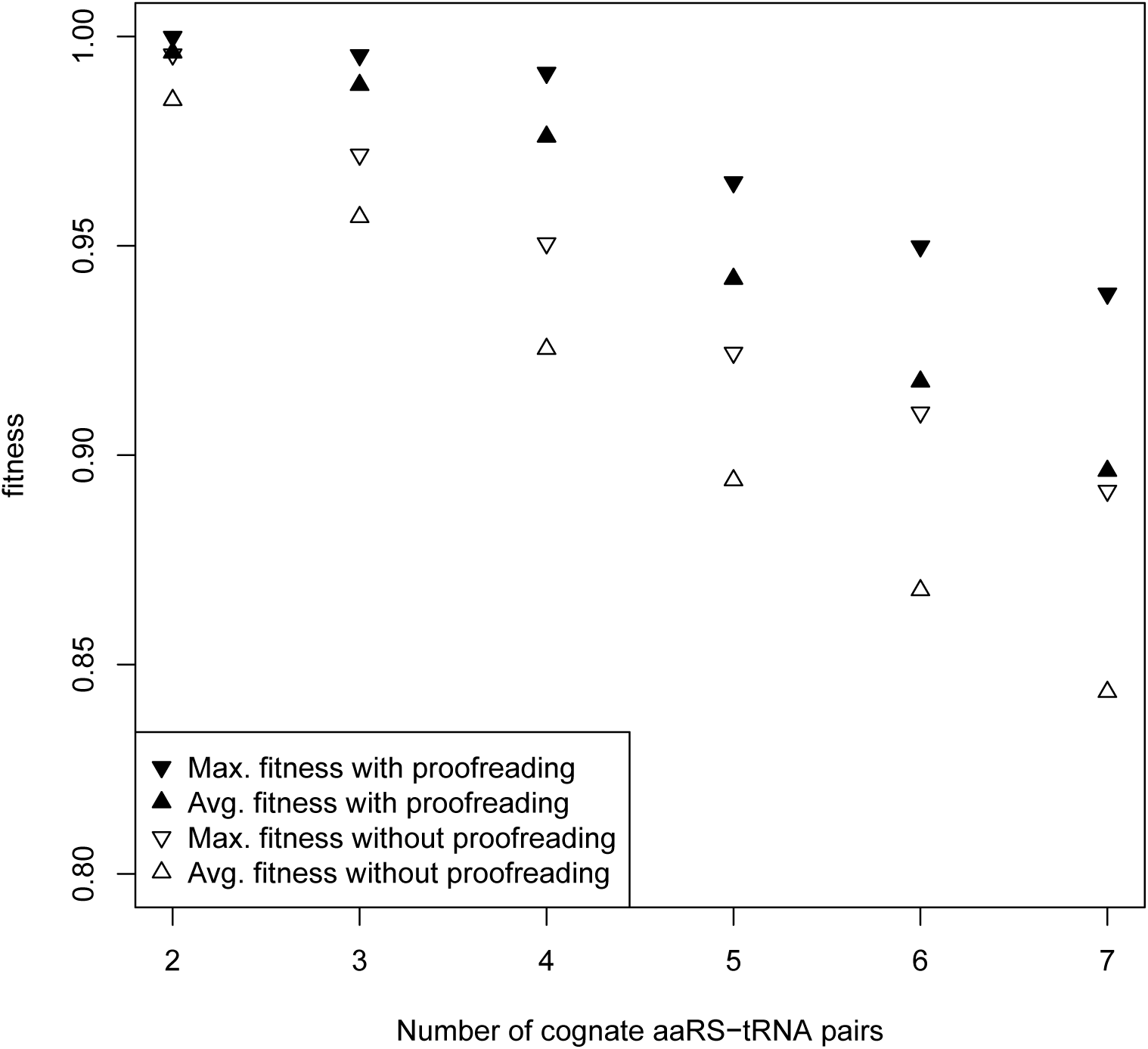
Decreasing average and maximal fitness of aaRS-tRNA networks as a function of encoded interactions under the XNOR rule, with and without proofreading. Parameters used here are *n* = 2, *k* = 2, *ϕ* = 0.9, and *β* = 100. The fixed-drift loads are the differences between maximal and average fitnesses, which increase with the number of encoded interactions. Notice the discontinuities between *P* = 4 and *P* = 5; this is the transition where *P* > 2^*n*^, the number of pairs exceeds available codewords. (one-column figure)

### 2.9. Selection on both translational accuracy and rate is necessary to select for masking to reduce cross-matching

The symmetric P-ary interaction channel as we have defined it, selects only for translational accuracy and not on rate or energy expenditure. One can see this clearly with the help of a well-defined example using the AND-XNOR rule, and comparing the fitnesses of two genotypes *g*_*H*_, *g*_*M*_ ∈ *G*^(4,4,8,2)^. The first genotype, *g*_*H*_, consists of four codewords from the Hamming [n=8,d=4] code [95] repeated twice, followed by all maskbits set:

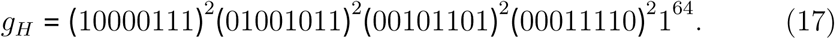

Since all maskbits are set in *g*_*H*_, all four tRNA species and all four aaRS species potentially match over their entire interfaces. The cognate match score for all pairs *t*_*i*_, *a*_*i*_ is *m*_*i*, *i*_ = 8 and the single non-cognate match score is *m*_*i,j*_ = 4 for all pairs *t*_*i*_, *a*_*j*_ with *i* ≠ *j*. No binary codes of size *n* = 8 can achieve a larger minimum Hamming distance than four [104, 105].

A second genotype *g*_*M*_ may be constructed from any two tetramers and their complements in the left and right halves of the interface, and using masking to eliminate cross-matching. For example,

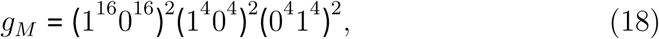

which achieves cognate matches (*m*_*i,i*_ = 4 for all pairs *t*_*i*_, *a*_*i*_) and zero crossmatching (*m*_*i,j*_ = 0 for all pairs *t*_*i*_, *a*_*j*_ with *i* ≠ *j*). With the standard fitness function that we have been using in which fitness depends only on accuracy and not rate of translation and using *k* = 4, the fitnesses of these two genotypes are exactly equal:

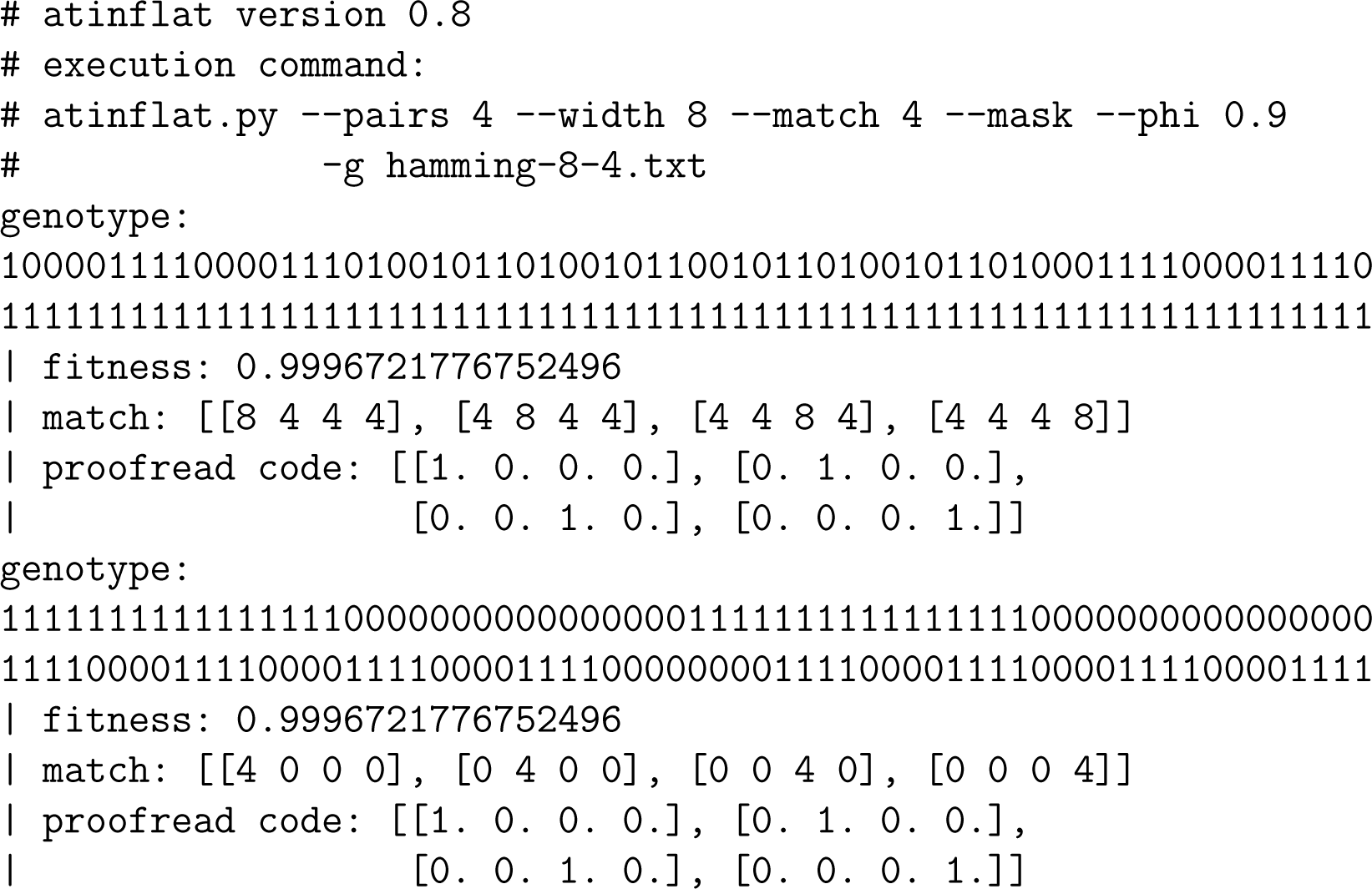

The example illustrates a key property of our macroscopic kinetic match landscape model, which is that accuracy depends on relative dissociation rate constants and concentrations, a prediction borne out by experimental evidence [34, 94, 106]. We conjecture that these two genotypes have maximal fitness because they both achieve the maximal possible distance of four between all code words — and they are not alone; many others in their neutral network have the same fitness. Other genotypes with equal fitness to *g*_*H*_ and *g*_M_ include all those with the structure of *g*_*H*_ but substituting any four of the 16 Hamming [8,4] codewords in any order, in any one of 2 × 8! permutations of codeword columns and codeword symbols (implying a degeneracy of more than 3.522 × 10^9^ Hamming code genotypes) as well as other non-linear perfect binary codes [107] — all with every mask-bit set — and a much smaller number of those with the same structure as *g*_*M*_ and half of the mask-bits off, using one of only 255 combinations of two tetramer codewords and their complements besides those used in *g*_*M*_.

Even though *g*_*H*_ and *g*_*M*_ achieve identical accuracy and fitness in the match landscape with *w*_*A*_(*g*), *g* ∈ *G*^(4,4,8,2)^, the rates of translation in cells with genotype *g*_*H*_ would be vastly slower than in cells with genotype *g*_*M*_, because the dissociation (discard) rate of cognate complexes is only between 1 s^−1^ and 2 s^−1^ in the former, while in the latter it is the typical cognate rate that we assumed, 220 s ^−1^. In the classic kinetic proofreading schemes, this discard rate must be much greater than the actual rate of product formation *k*_*cat*_ [56, 61] (but see [65, 66, 70]). For example, in tRNA-Ile of *Salmonella typhimurium* this rate is estimated to be 5 s ^−1^ [108]. Furthermore, the overall rate of protein synthesis, which factors directly into growth rate [109], can be limited by the slowest rate of aminoacylation [110, 111]. As a result, both the accuracy and rate of translation are expected to factor into fitness [112]. Because the fitnesses of *g*_*H*_ and *g*_*M*_ are exactly equal without taking translational rate into account, incorporating any rate-dependent fitness factor that decreases with the cognate aminoacylation rate in our model will disadvantage those genotypes that maximize matching between cognate complexes. Selection for accuracy should then select for mask bits to turn off to reduce cross-matching and maintain the high non-cognate/cognate dissociation rate ratios required for accuracy at intermediate levels of cognate matching.

To test this prediction, we introduce an empirically parametrized fitness factor that crudely penalizes cognate aminoacylation rates when they are slower than the assumed cognate rate of 220 s ^−1^. In accordance with an observation of *k*_cat_ = 5s ^−1^ [108] and a cognate dissociation/discard rate of 220 s^−1^, we define the average aminoacylation rate *k*_cat_(*g*) of genotype *g* as proportional to the harmonic mean of cognate dissociation rates between cognate tRNAs and aaRSs:

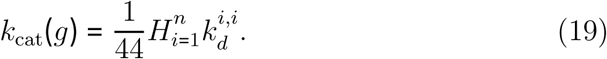

Controlled measurements with wild-type and mutant enzymes showed that only *k*_cat_ correlated with growth rate and the following measurements of (*k*_cat_, *w*) were observed, where *w* is growth rate in Luria Broth, written relative to wild-type [108, Table 3]: {(0.19, 0.24), (0.6, 0.6), (5,1)}.

Using GNUPLOT 5.2 to fit two exponential viability functions *w*_1_(*k*_cat_) = *A* + *B* exp(*C*_1_*k*_cat_) and *w*_2_(*k*_cat_) = 1-exp(*C*_2_*k*_cat_) to these data and also through the origin, we obtained the following fits:

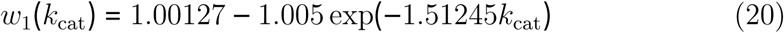

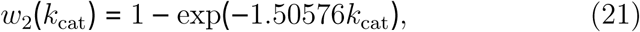

both with a root mean square residual of less than 1%.

We defined a new fitness function *w*_*AR*_(*g*) to select for both translational accuracy and rate as the product of two fitness factors:

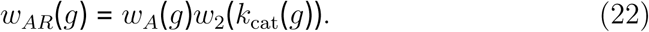

Using this new fitness function *w*_*AR*_(*g*) and *k* = 4, we obtained the following results:

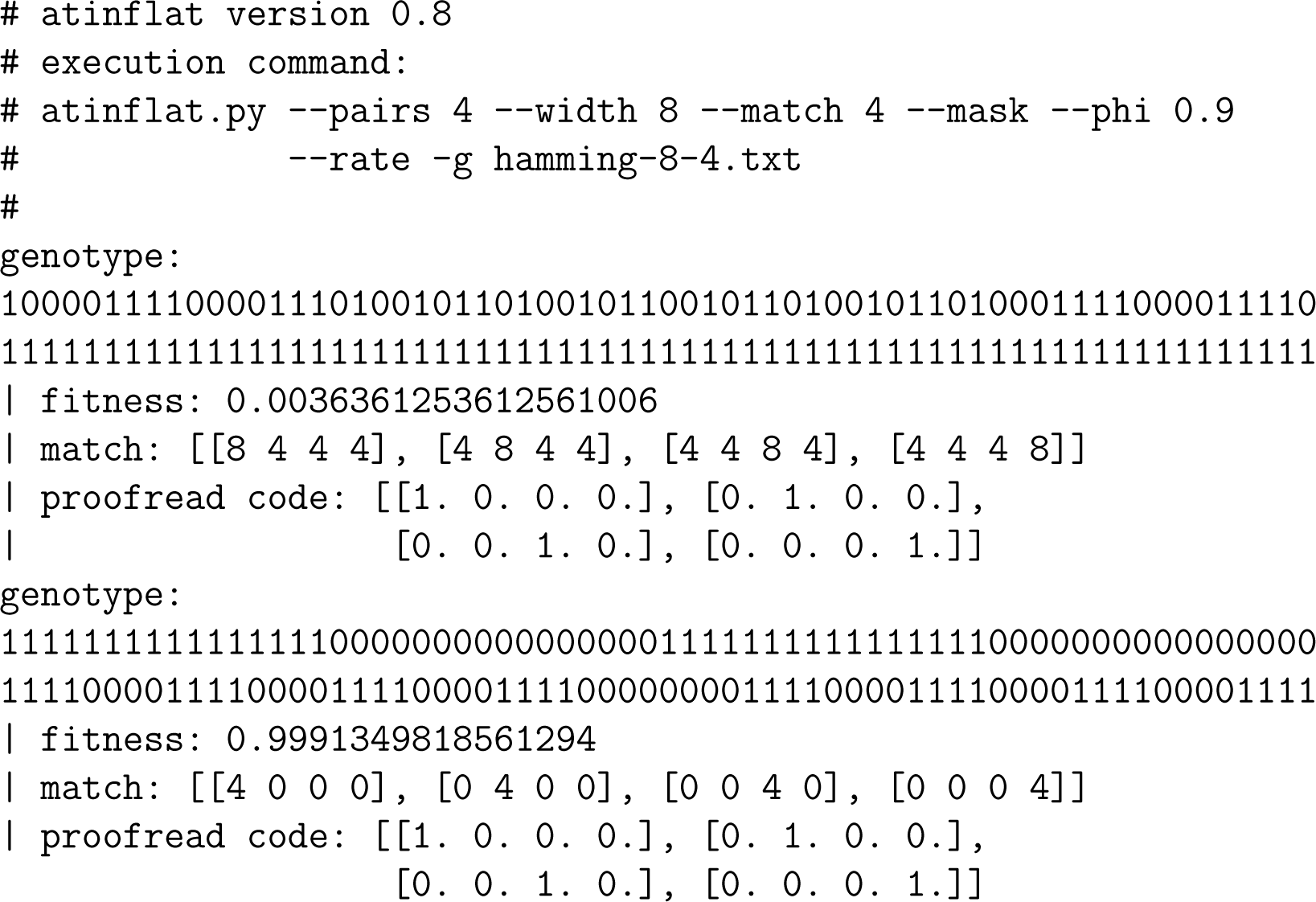

Even with *k* = 8, so the assumed cognate dissociation rate is only reached with a full eight matches, the masked genotype still has higher fitness:

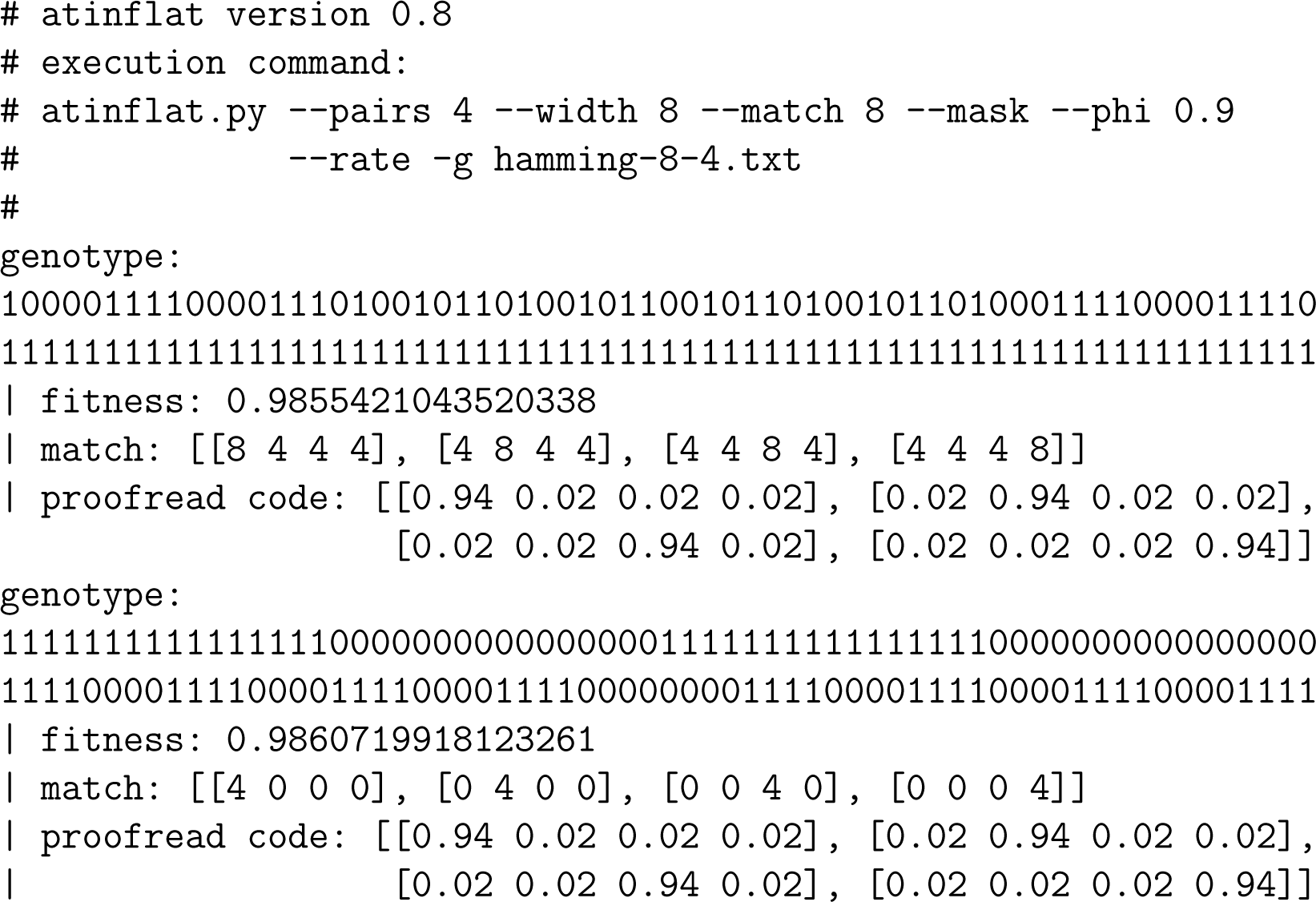

Hamming codes are efficient with respect to codeword length [95]. In this work, codewords are transmitted in parallel, so selection on code-word length *g*_*M*_ occurs through selection to avoid overly tight binding. Our results show that genetic match codes can be selected to sacrifice code-words to achieve shorter codeword length without cross-matching.

Our results are general. In Fig. 6, we show the full stationary genotype distributions under two fitness functions *w*_*A*_(*g*) and *w*_*AR*_(*g*) on the smaller genotype space *G*^(4,4,2,2)^ and *k* = 1, showing that masking is systematically favored over the entire match landscape and increasingly so with genotype fitness, under combined selection on translational accuracy and rate. Thus, selection on both the specificity of association and rate of dissociation can partition macromolecular interaction interfaces to reduce cross-matching.

**Figure 6:**
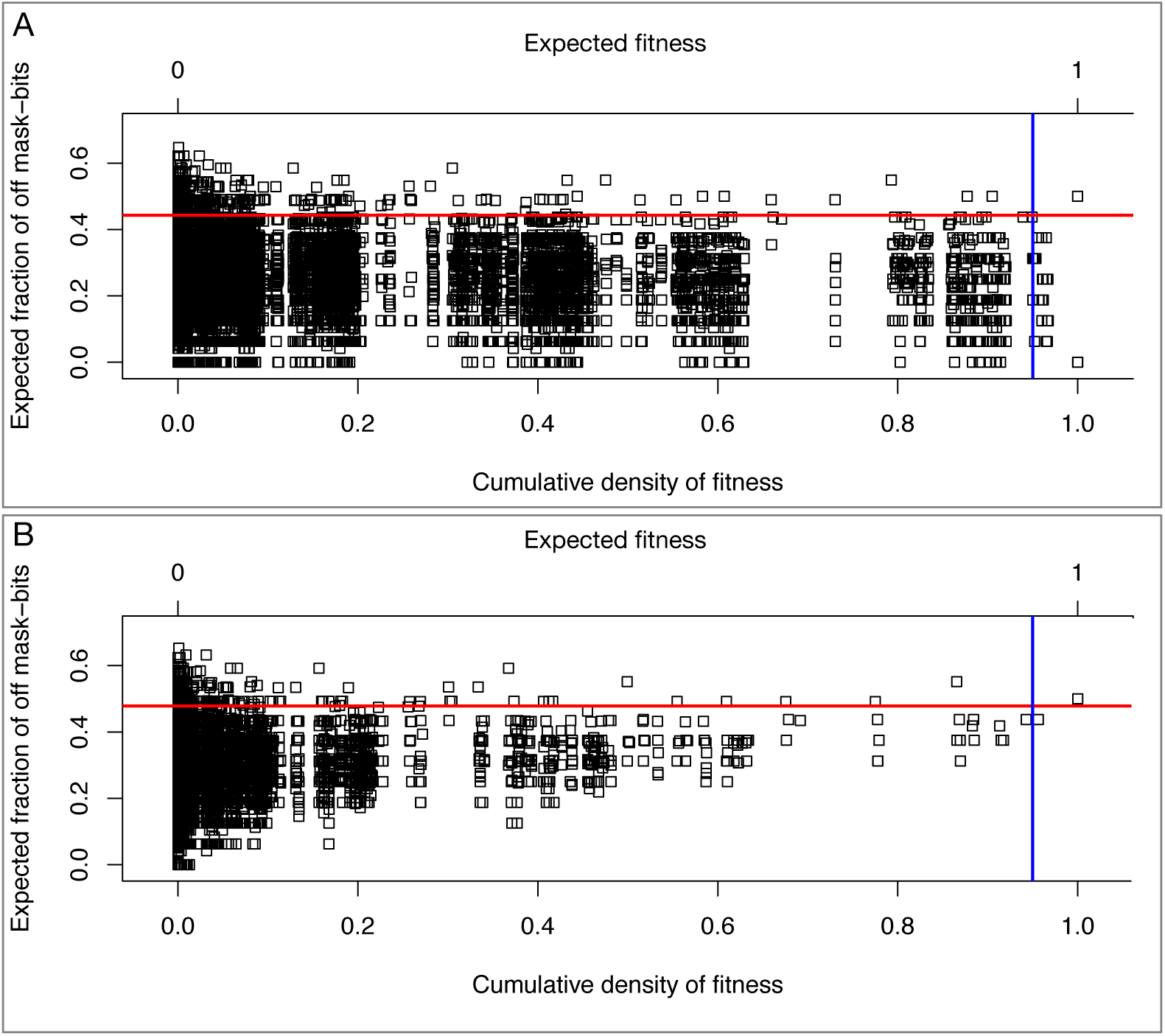
Expected fractions of masked sites in the steady-state fitness equivalence classes of 2^32^ genotypes in *G*^(4,4,2,2)^ (points), expected fractions of masked sites (red lines) and expected fitnesses (blue lines) as functions of the stationary cumulative densities of fitness in match landscapes with perfect one-step kinetic proofreading, *P* = 4, *n* = 2, *k* = 1, *ϕ* = 0.9, and *β* = 100. A. Match landscape with selection for translational accuracy alone (fitness function *w*_*A*_(*g*)) with expected fitness 0.9501304 and expected fraction of masked sites 0.4428638. B. Match landscape with combined selection for translational accuracy and rate (fitness function *w*_*AR*_(*g*)), with expected fitness is 0.9498575 and expected fraction of masked sites 0.4780706. Machine error in these data, as judged by the integration of cumulative density functions, is less than 10^−11^. (two-column figure)

Natural selection increases and maintains information in genomes [113–116]. A useful measure of this information is the reduction in entropy of the stationary distribution of genotypes with that selection, relative to without it. For example, the maximum entropy of genotypes in *G*^(4,4,2,2)^ occurs on a perfectly flat fitness landscape in which all genotypes have equal fitness, and its value is the genome length in bits, 32. For the data in Fig. 6 with perfect kinetic proofreading and *β* = 100, we found that the entropy of the stationary genotype distribution under selection for accuracy alone, through the fitness function *w*_*A*_(*g*), is about 7.82 bits for a maximum information gain of about 24.18 bits. The entropy of the stationary genotype distribution under combined selection for both accuracy and rate, through the fitness function *w*_*AR*_(*g*), is about 6.22 bits for a larger maximum information gain of about 25.78 bits. Thus, in a population of fixed size 101, about 1.6 more bits of information are gained under combined selection for rate and accuracy than under selection for accuracy alone. Without proofreading, the results are not very different: the maximum information gained under selection for accuracy alone is close to 23.5 bits, while under combined selection for both accuracy and rate, the maximum information gain is close to 25.26 bits.

## 3. Discussion

We have shown that combined selection on translational accuracy and rate is sufficient to select for divergence in tRNA-interaction interfaces by aaRSs. Our results do not contradict other hypotheses about this phenomenon [30]. We used mask bits as interaction modifiers to demonstrate our main result. When they mask or diminish interactions, these modifier bits may be interpreted as the presence of structural features such as *identity antideterminants* that prevent or weaken interactions at specific locations, possibly by guiding and orienting interactions away from other interaction-determining features [34].

The notion of “matching” used in this work should not be taken literally. The essential feature of the XNOR rule is its provision of two ways to match (0/0 and 1/1), corresponding to the availability of alternative paired sets of features in biomacromolecules that promote assortative interactions. As-sortative interactions occur by means of both *complementarity* in the shapes and motions of cognate pairs of tRNA and aaRS species, and *identifiability* or distinctiveness in the shapes and motions of cognate and non-cognate pairs. Because of the symmetry of mutation that we assumed in this work, we could have equivalently named our landscape a “complementarity landscape” and obtained identical results using an XOR matching rule instead of the XNOR rule. It would then be simple, although vague and misleading, to interpret matching features as complementarily charged amino acid side-chains or complementary RNA nucleobases that interact directly. However, this would be oversimplified on multiple levels: first, because identifying features in tRNAs can depend only indirectly on underlying bases and residues through the overall shape and motion in what is called *indirect read-out* [117]; second, tRNAs are extensively post-transcriptionally modified, which also biochemically integrates information from multiple sites in ways crucial for tRNA identity [118]; third, feature matching and mis-matching occurs in general through different sequence alphabets in RNA and proteins; and fourth, aaRS proteins are autocatalytically synthesized through the aaRS-tRNA network itself [119].

Thus, in the present work we analyzed only the simplest one of four increasingly complex variations on the general problem of evolution of a selfencoded aaRS-tRNA network. We define four connected notions to make our arguments: *description*, *self-description*, *self-encoded description*, and *self-encoded self-description*. By *description* we mean that when a mature, folded gene product evolves to complement the shape and motion of a fixed and unevolving ligand like a metabolite in order to specifically bind it, it “describes” that metabolite. This notion of “description” depends on the complex genotype-phenotype maps of RNA and protein folding, and therefore can attain complex and emergent evolutionary dynamics [91, 120]. Nonetheless, by definition, descriptions are of evolutionarily fixed targets and therefore intrinsically less rugged, with smaller neutral network size or degeneracy, than the match landscapes studied in the present work. We contend that evolving a description of an unevolvable metabolite ligand corresponds to discovering what might be called an *Easter egg* in sequence space. Under the assumption of symmetric mutation, the “all-ones” genotype studied in the Appendix of Sella [90] corresponds to selection to match any equivalent evolutionarily static Easter Egg in sequence space, of any arbitrary sequence neighborhood.

In the present work on the other hand, we analyzed the problem of *selfdescription*: specifically, we evolved co-inherited cognate tRNA-aaRS gene pairs to describe one another, so that their expressed products obtain complementary and identifying shapes and motions with one another. More generally, the notion of self-description represents the information acquired in genes by natural selection about the shapes and motions of the products (or regulatory regions) of other genes (which correspond to “self”’ with respect to the cell they are co-inherited in). During the evolutionary collectivization of genes and gene products into genomes and cells hypothesized by Woese and co-authors, genes acquired information via natural selection about the shapes and motions of other gene products, in order to interact specifically and/or conditionally with them. This *self-description* (or equivalently *self-information*) is the epistatic “biological glue” that binds folded macromolecules, cells and organisms together, enabling them to convert energy into work and execute complex emergent functions. Self-description applies equally well to epistatic interactions within genes and gene-products, where it programs their folds, major modes of motion, and allosteric changes in shape and motion in response to changes in cellular state. The reason that these notions of self-description are all consistent is precisely because the self-descriptions of biological entities at multiple scales become integrated through major evolutionary transitions.

Bedian [119] also called what he modeled “self-description,” but he meant something entirely different: the mutual self-compatible encoding of a set of aaRS catalytic active sites capable of aminoacylating different amino acids onto distinct tRNAs, so that the collection of self-encoded aaRSs active sites can autocatalytically resynthesize themselves and each other. In our terminological framework, this is *self-encoded description*, because tRNAs are treated as fixed and unevolving targets, like amino acids. Bedian’s model, and subsequent extensions by Wills and co-workers, consider that these different selectivities of different aaRSs depend on distinct sets of *critical sites* in each aaRS (where each critical site corresponds to one of our site-types). The distinct sets of critical sites of aaRSs may be thought of as multiple distinct Easter eggs in sequence space that all must be simultaneously discovered and compatibly mutually encoded for the network of aaRS active sites to nucleate. But aaRSs have both catalytic and tRNA-binding domains. Bedian, Wills and co-workers have so far not considered the problem of tRNA recognition by autocatalytically encoded aaRSs in their work, which generalizes what we studied here in what might be called *self-encoded self-description*. Full treatment of the problem, involving autocatalytically-encoded Easter eggs and Match Landscapes, is reserved for future investigations. Progress will allow a fuller investigation of even larger models to investigate the coevolution of genetic code and metabolism [121, 122].

We conjecture that our present results will hold for these more complex models. We offer an interpretation of “matching” for our present results which applies to all of these more complex biological settings; namely, matching represents the self-information contained in self-descriptions, or the information contained in genes about the identifying shapes and motions of other co-inherited genes and gene products. Commensurately informative self-descriptions are expected to be *nearly neutral* with one another in the sense of [90] and references therein, and as shown for interaction interfaces previously [123]. The nearly-neutral evolution of interaction-determining features within a high-dimensional sequence space of equally fit solutions makes compensatory mutations much more likely than reversals. This explains both why interaction-informative features can evolve and diverge even while under strong selection, and why interaction-determining features are phylogeneti-cally informative.

Our theory that macromolecular interactions are encoded through sets of complementary and identifying features extends the universal principle of heredity clarified by Watson and Crick, through which all possible genetic sequences may be replicated by virtue of complementarity [124]. The relativity of the notions of complementarity and identity in the definition of self-description implies that macromolecular interactions are governed by symbolic representations, as discussed by Maynard Smith [115]. That is, within the context of a specific cell, arbitrary molecular shapes and motions are symbolically associated with specific functions. The notion of symbolic association is defined not only by the absence of relationship between the form and meaning of signals [115], but also by its cryptographic nature, in that it requires coordinated information to decode signals correctly within a large space of equally unambiguously expressive alternatives.

The statistical Tree of Life became more tree-like after the Darwinian Transition precisely because through this transition, cells evolved languages of self-encoded descriptions and self-descriptions critical to their fitness as cells. These genetic and cellular languages are symbolic, crytographic, open-endedly expressive, and increasingly constrained from changing by the increasingly complex corpus of descriptions and self-descriptions they encode. Since languages evolve in a statistically tree-like manner [125, 126], so did the advent of these cellular and genetic languages caused cells to evolve in a statistically tree-like manner. Furthermore, the large degeneracy of equivalent self-descriptions implies that such a language may be surprisingly easy to originate spontaneously, yet once originated, will be heavily constrained to change only in ancestrally compatible ways [12].

It is easy to imagine that macromolecular interaction codes, like languages, evolve to be both expressive and unambiguous, that is, to encode more and more interactions in robust and error-tolerant (and ambiguity-reducing) ways. The coding theory analogy to the universality of replication by complementarity lies in the notion of *non-trivial perfect codes*. Perfect codes uniquely cover all of a finite sequence space with a maximum number of code-words spaced a minimum distance apart, so that every single possible code-word can be received unambiguously and decoded correctly even after one or more symbols in the code-word were altered. While we expect biological codes to be generally far from perfect, the theory of perfect codes may be a useful reference point from which to relax assumptions, and seems relevant to the stochastic setting of gene expression. In this context, it is of interest to note that surprisingly few varieties of small non-trivial perfect codes exist (where *non-trivial* means a code with more than one code word, not using every possible sequence as a codeword, nor the *P*-ary repetition code) [102]. For symbolic alphabets of prime power size, all non-trivial perfect codes have codeword sizes, lengths, and minimum distance parameters equal to those of either Hamming Codes or Golay Codes [102, 127]. However, the Golay codes are too large to be relevant to the problem of perfect coding of 20 or fewer aaRS-tRNA cognate interactions. The RNA alphabet is of prime power size, namely four. The Hamming code ℋ_*r*_(*h*) over an alphabet of size *r* with positive integer index parameter *h* has *M* = *r*^*n*-*h*^ codewords of length *n* = (*r*^*h*^ − 1)/(*r* − 1) and minimum Hamming distance between codewords of 3, allowing correction of single-symbol errors. It is of interest to note that ℋ_4_(2) contains four codewords of size 5, ℋ_4_(3) contains 16 codewords of size 21, and ℋ_4_(4) contains 64 codewords of size 85. The ℋ_4_(3) perfect codeword length of 21 is surprisingly close to the size of a postulated primordial tRNA hairpin [24, 128, 129] with acceptor stem length of 7 and anticodon loop of length 7, while the ℋ_4_(4) perfect codeword length of 85 is surprisingly close to the typical lengths of tRNAs today.

We can use our theory to roughly calculate the probability *p*(*n*, *P*, *d*, *H*) that an aaRS-tRNA network will evolve *P* matching codewords of minimum distance *d* over an interface of length *n* in a system with *M* mutually dissimilar tRNA replicators and *N* mutually dissimilar aaRS ribozyme replicators (with *P* ≤ *M*, *N*), and aaRS-tRNA per-site background and target symbol-pair frequencies defined by the expected relative entropy *H*. Counting all possible pairs between tRNA and aaRS genes, and assuming that tRNAs have evolveable anticodons, this probability is

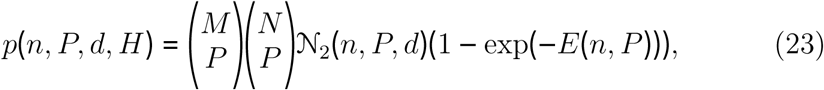

where 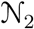(*n*, *P*, *d*) is the number of binary codes of length *n*, size *P* and minimum Hamming distance *d*, *E*(*n*, *P*) = *kMNn*^2^2^-*nPH*^ is the expected number of random sequences achieving normalized score *nPH* in a search space of size *nM* × *nN*, from Karlin-Altschul theory (and in which *k* is a correction factor for edge effects) [130], and where the expected relative entropy per-site *H* may be computed by enumerating over all pairs of RNA bases, assuming a specific base composition common to all genes, and an expected target similarity corresponding to one or fewer errors per *n* symbols. Although a unique finite number of codes 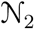 (*n*, *P*, *d*) exists over any finite sequence space, no expression for its value is known [127]. However this number must be much larger than the number of ways to choose *P* codewords from any Hamming-code of length *n* and size *Q* ≥ *P*, provided Hamming codes of that length and size exist, because of the existence of a potentially large, yet unknown number of non-linear codes with Hamming parameters [127]. The number of distinct Hamming codes of length *n* over an alphabet of size *q* is *q*!*n*! [127]. Further investigation is needed, but we believe that *p*(*n*, *P*, *d*, *H*) may be surprisingly large.

The theory by which we computed stationary genotype distributions can incorporate up to three kinds of mutational asymmetry [88] such as GC-bias, transition bias, or transcription-or strand-dependent mutation, all relevant to problems in the evolution of the genetic code. It should be expected that incorporating asymmetric mutation will break symmetries in the fitness of genotypes and will change the expected composition of interaction-determining features.

An importantly unrealistic assumption in the present work is that of large aaRS concentrations in our macroscopic model of aminoacylation. The stochastic dynamics of cellular-scale aminoacylation coupled to the sink of translating ribosomes is complex, exhibiting phenomena such as ultra-sensitivity [111]. We have implemented a mesoscopic version of aminoacylation kinetics using Gillespie’s direct method [131], results with which will be published elsewhere. Although our results do not depend on how translational rate is implemented, our model can fruitfully be integrated into a fully stochastic model of translation such as in Shah et al. [132]. In future work we will incorporate these and other extensions into new models for the coevolution of genetically encoded descriptions and self-descriptions with codon meanings and metabolism in structured populations, to better understand evolution through the Darwinian Transition.

## 4. Acknowledgments

DHA and AC-H were supported by the National Science Foundation (INSPIRE-1344279). DHA was supported by a Julius Kuhn Guest Professorship at Martin Luther Universitat in Halle-Wittenberg. Computational research in this report was performed on the MERCED HPC cluster sup-ported by the National Science Foundation (ACI-1429783). Our funding sources had no role in study design, analysis, interpretation, writing or submission. We thank Harish Bhat, Ivo Grosse, Suzanne Sindi, Kyle Kauffman, Michael Frisch, and Emily Jane McTavish for valuable discussions.

## Appendix A. Decomposition of the steady state solution of fixed genotypes with multiplicative fitness components

**Remark 1**. Define 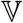 to be the set of possible values in a site. 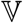 could be the set of nucleotides, the set of amino acids, etc. In this particular study, 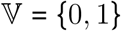

**Remark 2**. Define G^(*M,N,n,p*)^ (*M* not necessarily unequal to *N*) to be the set of all possible genotypes of width *n* ∈ ℕ and *pn* sites per-gene, with *p* ∈ ℕ. If ᗄ*g* ∈ *G*^(*M,N,n,p*)^ have length *L*, then 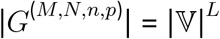.

**Remark 3**. Consider a genotype, *g* ∈ *G*^(*M,N,n,p*)^. Let 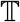 be the set of tRNA genes in 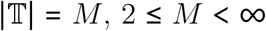 and let 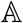 be the set of aaRS genes in *g* with 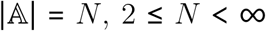. The lengths of genes 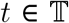 and 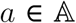 are all equal to *np* ᗄ*t*, α. Let *p*(*M* + *N*) = *L_b_*. Define **block** 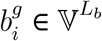, *i* ∈ {1, 2,…, *n*} to be the sequence of *p* ordered values starting at the *j*^*th*^ site across all *t* and *a* genes in genotype *g*, with *j* = (*i* − 1)*p*. For a genotype *g* ∈ *G*^(*M,N,n,p*)^, there will be *n* blocks 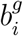, and each will be *L*_*b*_ long, it is possible for 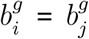 for 1 ≤ *i* ≠ *j* ≤ *n*, and 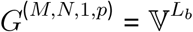 is the set of all possible types of blocks.

**Theorem 1**. *Let w*_*g*_ *be the viability of genotype* 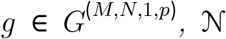 *be the population size, and* 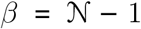 *for the Moran process*, 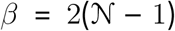 *for the haploid Wright-Fisher process, and* 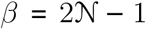 *for the diploid Wright-Fisher process. Given that the stationary frequency* 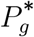 *of genotype g is* 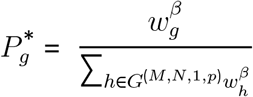 *and that the viability W*_κ_ *is multiplicative across blocks in a genotype κ* ∈ *G*^(*M,N,n*>1, *p*)^ 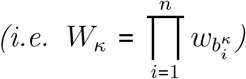 *then the stationary frequency* 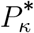 *of genotype* κ *is*

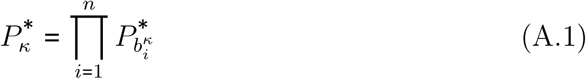

Proof (Proof of A.1). 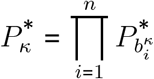

By definition,

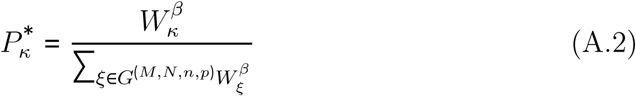

By the multiplicativity property this becomes

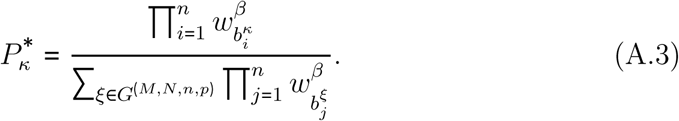

It needs to be shown that 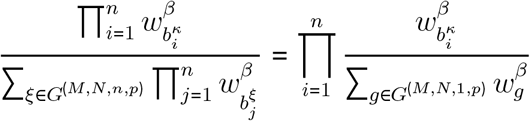 Essentially, the proof breaks down to whether 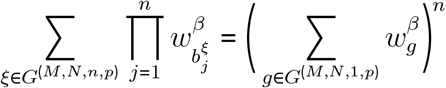 Start with,

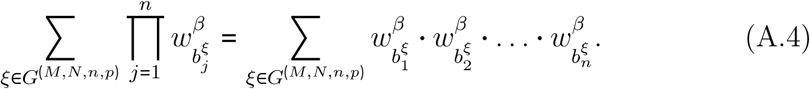

Since *G*^(*M,N*,1, *p*)^ is the set of all possible blocks, 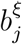, and no combination of *L*_*b*_ length genotypes across blocks is impossible, there are *B*^*n*^ possible sequences for genotypes ξ ∈ *G*^(*M,N,n,p*)^, where *B* = |*G*^(*M,N*,1, *p*)^|. This is consistent with the cardinality of *G*^(*M,N,n,p*)^ since *L* = *L*_*b*_*n* and thus 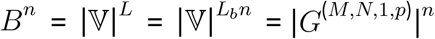. Since we are summing over all possible genotypes ξ ∈ *G*^(*M,N,n,p*)^, and since different genotypes in *G*^(*M,N,n,p*)^ with the same blocks but in different orders will have the same viability, then every viability term will be of the form 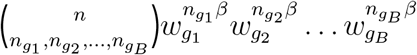 where each *g*_*i*_ ∈ *G*^(*M,N*,1, *p*)^ is (possibly arbitrarily) ordered from 1 to *B* and 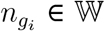 is the number of blocks of genotype ξ that are *g_i_*. Since every genotype is represented, (A.4) is a multinomial and can be rewritten 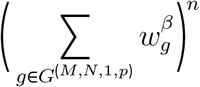 If this were not the case and one of the viability coefficients was less than the expected multinomial coefficient, then that could only mean that at least one genotype was not being counted. If one had a coefficient larger than expected it would have to mean that at least one genotype was being counted more than once. Therefore to prove (A.1), plug this multinomial representation into (A.3),

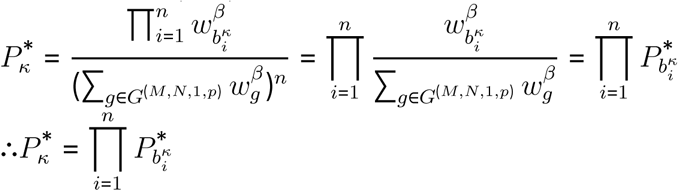

## References

[1] C. R. Woese, Interpreting the universal phylogenetic tree, Proceedings of the National Academy of Sciences 97 (2000) 8392–8396.

[2] C. R. Woese, G. J. Olsen, M. Ibba, D. Söll, Aminoacyl-tRNA synthetases, the genetic code, and the evolutionary process, Microbiology and molecular biology reviews: MMBR 64 (2000) 202–236.

[3] C. R. Woese, On the evolution of cells, Proceedings of the National Academy of Sciences 99 (2002) 8742–8747.

[4] K. Vetsigian, C. Woese, N. Goldenfeld, Collective evolution and the genetic code, Proceedings of the National Academy of Sciences 103 (2006) 10696–10701.

[5] E. Roberts, A. Sethi, J. Montoya, C. R. Woese, Z. Luthey-Schulten, Molecular signatures of ribosomal evolution, Proceedings of the National Academy of Sciences of the United States of America 105 (2008) 13953–13958.

[6] T. Ruusala, D. Andersson, M. Ehrenberg, C. G. Kurland, Hyper-accurate ribosomes inhibit growth., The EMBO Journal 3 (1984) 2575–2580.

[7] C. G. Kurland, Translational accuracy and the fitness of bacteria, Annual Review of Genetics 26 (1992) 29–50.

[8] W. Ran, P. G. Higgs, Contributions of Speed and Accuracy to Translational Selection in Bacteria, PLOS ONE 7 (2012) e51652.

[9] L. Ribas de Pouplana, M. A. S. Santos, J.-H. Zhu, P. J. Farabaugh, B. Javid, Protein mistranslation: friend or foe?, Trends in Biochemical Sciences 39 (2014) 355–362.

[10] E. V. Koonin, The Biological Big Bang model for the major transitions in evolution, Biology Direct 2 (2007) 21.

[11] M. L. Katz, C. Shapiro, Systems Competition and Network Effects, The Journal of Economic Perspectives 8 (1994) 93–115.

[12] F. H. Crick, The origin of the genetic code, Journal of Molecular Biology 38 (1968) 367–379.

[13] E. Szathmry, J. M. Smith, The major evolutionary transitions, Nature 374 (1995) 227–232.

[14] M. M. Desai, D. Weissman, M. W. Feldman, Evolution Can Favor Antagonistic Epistasis, Genetics 177 (2007) 1001–1010.

[15] U. Liberman, M. Feldman, On the evolution of epistasis III: the haploid case with mutation, Theoretical Population Biology 73 (2008) 307–316.

[16] E. V. Koonin, Horizontal gene transfer: essentiality and evolvability in prokaryotes, and roles in evolutionary transitions, F1000Research 5 (2016).

[17] M. C. Rivera, R. Jain, J. E. Moore, J. A. Lake, Genomic evidence for two functionally distinct gene classes, Proceedings of the National Academy of Sciences of the United States of America 95 (1998) 6239–6244.

[18] R. Jain, M. C. Rivera, J. A. Lake, Horizontal gene transfer among genomes: the complexity hypothesis, Proceedings of the National Academy of Sciences of the United States of America 96 (1999) 3801–3806.

[19] H. B. Fraser, A. E. Hirsh, L. M. Steinmetz, C. Scharfe, M. W. Feldman, Evolutionary rate in the protein interaction network, Science (New York, N.Y.) 296 (2002) 750–752.

[20] C. S. Francklyn, E. A. First, J. J. Perona, Y.-M. Hou, Methods for kinetic and thermodynamic analysis of aminoacyl-tRNA synthetases, Methods (San Diego, Calif.) 44 (2008) 100–118.

[21] C. Francklyn, K. Musier-Forsyth, S. A. Martinis, Aminoacyl-tRNA synthetases in biology and disease: new evidence for structural and functional diversity in an ancient family of enzymes, RNA (New York, N.Y.) 3 (1997) 954–960.

[22] Y. I. Wolf, L. Aravind, N. V. Grishin, E. V. Koonin, Evolution of aminoacyl-tRNA synthetases-analysis of unique domain architectures and phylogenetic trees reveals a complex history of horizontal gene transfer events, Genome Research 9 (1999) 689–710.

[23] P. O’Donoghue, Z. Luthey-Schulten, On the evolution of structure in aminoacyl-tRNA synthetases, Microbiology and molecular biology reviews: MMBR 67 (2003) 550–573.

[24] P. Schimmel, R. Giege, D. Moras, S. Yokoyama, An operational RNA code for amino acids and possible relationship to genetic code, Proceedings of the National Academy of Sciences of the United States of America 90 (1993) 8763–8768.

[25] S. Shaul, D. Berel, Y. Benjamini, D. Graur, Revisiting the operational RNA code for amino acids: Ensemble attributes and their implications, RNA (New York, N.Y.) 16 (2010) 141–153.

[26] G. Eriani, M. Delarue, O. Poch, J. Gangloff, D. Moras, Partition of tRNA synthetases into two classes based on mutually exclusive sets of sequence motifs, Nature 347 (1990) 203–206.

[27] S. Cusack, Aminoacyl-tRNA synthetases, Current Opinion in Structural Biology 7 (1997) 881–889.

[28] J. R. Brown, W. F. Doolittle, Root of the universal tree of life based on ancient aminoacyl-tRNA synthetase gene duplications, Proceedings of the National Academy of Sciences of the United States of America 92 (1995) 2441–2445.

[29] L. Ribas de Pouplana, P. Schimmel, Two classes of tRNA synthetases suggested by sterically compatible dockings on tRNA acceptor stem, Cell 104 (2001) 191–193.

[30] L. Ribas de Pouplana, P. Schimmel, Aminoacyl-tRNA synthetases: potential markers of genetic code development, Trends in Biochemical Sciences 26 (2001) 591–596.

[31] D. S. Goodsell, S. Dutta, C. Zardecki, M. Voigt, H. M. Berman, S. K. Burley, The RCSB PDB Molecule of the Month: Inspiring a Molecular View of Biology, PLOS Biology 13 (2015) e1002140.

[32] M. Sprinzl, F. Grueter, A. Spelzhaus, D. H. Gauss, Compilation of tRNA sequences., Nucleic Acids Research 8 (1980) r1–r22.

[33] R. Giege, E. Touze, B. Lorber, A. Theobald-Dietrich, C. Sauter, Crystallogenesis Trends of Free and Liganded Aminoacyl-tRNA Synthetases, Crystal Growth & Design 8 (2008) 4297–4306.

[34] R. Giege, M. Sissler, C. Florentz, Universal rules and idiosyncratic features in tRNA identity, Nucleic Acids Research 26 (1998) 5017–5035.

[35] E. Freyhult, V. Moulton, D. H. Ardell, Visualizing bacterial tRNA identity determinants and antideterminants using function logos and inverse function logos, Nucleic Acids Research 34 (2006) 905–916.

[36] D. H. Ardell, S. G. E. Andersson, TFAM detects co-evolution of tRNA identity rules with lateral transfer of histidyl-tRNA synthetase, Nucleic Acids Research 34 (2006) 893–904.

[37] D. H. Ardell, G. Sella, On the evolution of redundancy in genetic codes, Journal of Molecular Evolution 53 (2001) 269–281.

[38] E. Freyhult, Y. Cui, O. Nilsson, D. H. Ardell, New computational methods reveal tRNA identity element divergence between Proteobac-teria and Cyanobacteria, Biochimie 89 (2007) 1276–1288.

[39] K. C. H. Amrine, W. D. Swingley, D. H. Ardell, tRNA signatures reveal a polyphyletic origin of SAR11 strains among alphaproteobacteria, PLoS computational biology 10 (2014) e1003454.

[40] E. V. Koonin, A. S. Novozhilov, Origin and evolution of the genetic code: the universal enigma, Iubmb Life 61 (2009) 99–111.

[41] S. E. Massey, The neutral emergence of error minimized genetic codes superior to the standard genetic code, Journal of Theoretical Biology 408 (2016) 237–242.

[42] S. N. Rodin, S. Ohno, Two types of aminoacyl-trna synthetases could be originally encoded by complementary strands of the same nucleic ACID, Origins of life and evolution of the biosphere 25 (1995) 565–589.

[43] S. N. Rodin, A. S. Rodin, On the origin of the genetic code: signatures of its primordial complementarity in tRNAs and aminoacyl-tRNA synthetases, Heredity 100 (2008) 341–355.

[44] C. W. Carter, L. Li, V. Weinreb, M. Collier, K. Gonzalez-Rivera, M. Jimenez-Rodriguez, O. Erdogan, B. Kuhlman, X. Ambroggio, T. Williams, S. N. Chandrasekharan, The Rodin-Ohno hypothesis that two enzyme superfamilies descended from one ancestral gene: an unlikely scenario for the origins of translation that will not be dismissed, Biology Direct 9 (2014) 11.

[45] J. A. G. M. de Visser, J. Hermisson, G. P. Wagner, L. Ancel Meyers, H. Bagheri-Chaichian, J. L. Blanchard, L. Chao, J. M. Cheverud, S. F. Elena, W. Fontana, G. Gibson, T. F. Hansen, D. Krakauer, R. C. Lewontin, C. Ofria, S. H. Rice, G. von Dassow, A. Wagner, M. C. Whitlock, Perspective: Evolution and detection of genetic robustness, Evolution; International Journal of Organic Evolution 57 (2003) 1959–1972.

[46] C. O. Wilke, J. L. Wang, C. Ofria, R. E. Lenski, C. Adami, Evolution of digital organisms at high mutation rates leads to survival of the flattest, Nature 412 (2001) 331–333.

[47] O. G. Berg, P. H. von Hippel, Selection of DNA binding sites by regulatory proteins. Statistical-mechanical theory and application to operators and promoters, Journal of Molecular Biology 193 (1987) 723–750.

[48] U. Gerland, J. D. Moroz, T. Hwa, Physical constraints and functional characteristics of transcription factor-DNA interaction, Proceedings of the National Academy of Sciences of the United States of America 99 (2002) 12015–12020.

[49] J. Berg, M. Lössig, A. Wagner, Structure and evolution of protein interaction networks: a statistical model for link dynamics and gene duplications, BMC Evolutionary Biology 4 (2004) 51.

[50] V. Mustonen, J. Kinney, C. G. Callan, M. Lassig, Energy-dependent fitness: A quantitative model for the evolution of yeast transcription factor binding sites, Proceedings of the National Academy of Sciences of the United States of America 105 (2008) 12376–12381.

[51] A. Y. Tulchinsky, N. A. Johnson, W. B. Watt, A. H. Porter, Hybrid incompatibility arises in a sequence-based bioenergetic model of transcription factor binding, Genetics 198 (2014) 1155–1166.

[52] T. Friedlander, R. Prizak, C. C. Guet, N. H. Barton, G. Tkacik, Intrinsic limits to gene regulation by global crosstalk, Nature Communications 7 (2016) 12307.

[53] E. J. Deeds, O. Ashenberg, J. Gerardin, E. I. Shakhnovich, Robust pro-teinprotein interactions in crowded cellular environments, Proceedings of the National Academy of Sciences 104 (2007) 14952–14957.

[54] M. E. Johnson, G. Hummer, Nonspecific binding limits the number of proteins in a cell and shapes their interaction networks, Proceedings of the National Academy of Sciences of the United States of America 108 (2011) 603–608.

[55] M. E. Johnson, G. Hummer, Interface-Resolved Network of Protein-Protein Interactions, PLoS Computational Biology 9 (2013).

[56] J. J. Hopfield, Kinetic Proofreading: A New Mechanism for Reducing Errors in Biosynthetic Processes Requiring High Specificity, Proceedings of the National Academy of Sciences 71 (1974) 4135–4139.

[57] J. Ninio, Kinetic amplification of enzyme discrimination, Biochimie 57 (1975) 587–595.

[58] K.-W. Leong, U. Uzun, M. Selmer, M. Ehrenberg, Two proofreading steps amplify the accuracy of genetic code translation, Proceedings of the National Academy of Sciences of the United States of America 113 (2016) 13744–13749.

[59] T. Yamane, J. J. Hopfield, Experimental evidence for kinetic proofreading in the aminoacylation of tRNA by synthetase., Proceedings of the National Academy of Sciences of the United States of America 74 (1977) 2246–2250.

[60] H. Mellenius, M. Ehrenberg, Transcriptional accuracy modeling suggests two-step proofreading by RNA polymerase, Nucleic Acids Research 45 (2017) 11582–11593.

[61] H. Qian, Reducing Intrinsic Biochemical Noise in Cells and Its Thermodynamic Limit, Journal of Molecular Biology 362 (2006) 387–392.

[62] W. S. Hlavacek, A. Redondo, H. Metzger, C. Wofsy, B. Goldstein, Kinetic proofreading models for cell signaling predict ways to escape kinetic proofreading, Proceedings of the National Academy of Sciences of the United States of America 98 (2001) 7295–7300.

[63] T. W. McKeithan, Kinetic proofreading in T-cell receptor signal transduction, Proceedings of the National Academy of Sciences 92 (1995) 5042–5046.

[64] V. Barone, M. Lang, S. F. G. Krens, S. J. Pradhan, S. Shamipour, K. Sako, M. Sikora, C. C. Guet, C.-P. Heisenberg, An Effective Feedback Loop between Cell-Cell Contact Duration and Morphogen Signaling Determines Cell Fate, Developmental Cell 43 (2017) 198–211.e12.

[65] A. Murugan, D. A. Huse, S. Leibler, Speed, dissipation, and error in kinetic proofreading, Proceedings of the National Academy of Sciences 109 (2012) 12034–12039.

[66] K. Banerjee, A. B. Kolomeisky, O. A. Igoshin, Elucidating interplay of speed and accuracy in biological error correction, Proceedings of the National Academy of Sciences of the United States of America 114 (2017) 5183–5188.

[67] M. Ehrenberg, C. Blomberg, Thermodynamic constraints on kinetic proofreading in biosynthetic pathways, Biophysical Journal 31 (1980) 333–358.

[68] C. G. Kurland, The role of guanine nucleotides in protein biosynthesis., Biophysical Journal 22 (1978) 373–392.

[69] C.-M. Zhang, J. J. Perona, K. Ryu, C. Francklyn, Y.-M. Hou, Distinct kinetic mechanisms of the two classes of Aminoacyl-tRNA synthetases, Journal of Molecular Biology 361 (2006) 300–311.

[70] P. Sartori, S. Pigolotti, Kinetic versus Energetic Discrimination in Biological Copying, Physical Review Letters 110 (2013) 188101.

[71] Wright, S., The roles of mutation, inbreeding, crossbreeding and selection in evolution., Proc. 6th Int. Congress on Genetics, Ithaca, NY, USA 1 (1932) 356–366.

[72] S. E. Ahnert, Structural properties of genotypephenotype maps, Journal of the Royal Society Interface 14 (2017).

[73] Dykhuizen, D, Recommendation of [Lunzer M et al., Science 2005, 310(5747):499–501], 2005.

[74] K. Crona, D. Greene, M. Barlow, The peaks and geometry of fitness landscapes, Journal of Theoretical Biology 317 (2013) 1–10.

[75] F. J. Poelwijk, S. Tnase-Nicola, D. J. Kiviet, S. J. Tans, Reciprocal sign epistasis is a necessary condition for multi-peaked fitness landscapes, Journal of Theoretical Biology 272 (2011) 141–144.

[76] Kauffman, S.A., The Origins of Order. Self-Organization and Selection in Evolution., Oxford University Press, Oxford, U.K., 1993.

[77] S. Kauffman, S. Levin, Towards a general theory of adaptive walks on rugged landscapes, Journal of Theoretical Biology 128 (1987) 11–45.

[78] M. L. Siegal, A. Bergman, Waddington’s canalization revisited: Developmental stability and evolution, Proceedings of the National Academy of Sciences 99 (2002) 10528–10532.

[79] T. MacCarthy, A. Bergman, Coevolution of robustness, epistasis, and recombination favors asexual reproduction, Proceedings of the National Academy of Sciences 104 (2007) 12801–12806.

[80] A. Orlenko, P. B. Chi, D. A. Liberles, Characterizing the roles of changing population size and selection on the evolution of flux control in metabolic pathways, BMC evolutionary biology 17 (2017) 117.

[81] M. Nei, Modification of Linkage Intensity by Natural Selection, Genetics 57 (1967) 625–641.

[82] Feldman, M. W., Selection for linkage modification: I. Random mating populations., Theoretical Population Biology 3 (1972) 324–346.

[83] L. Altenberg, U. Liberman, M. W. Feldman, Unified reduction principle for the evolution of mutation, migration, and recombination, Proceedings of the National Academy of Sciences 114 (2017) E2392–E2400.

[84] C. O. Wilke, C. Adami, Interaction between directional epistasis and average mutational effects, Proceedings of the Royal Society of London B: Biological Sciences 268 (2001) 1469–1474.

[85] P.-A. Gros, H. L. Nagard, O. Tenaillon, The Evolution of Epistasis and Its Links With Genetic Robustness, Complexity and Drift in a Phenotypic Model of Adaptation, Genetics 182 (2009) 277–293.

[86] D. M. Weinreich, R. A. Watson, L. Chao, Perspective: Sign epistasis and genetic constraint on evolutionary trajectories, Evolution; International Journal of Organic Evolution 59 (2005) 1165–1174.

[87] D. B. Weissman, M. M. Desai, D. S. Fisher, M. W. Feldman, The rate at which asexual populations cross fitness valleys, Theoretical Population Biology 75 (2009) 286–300.

[88] G. Sella, A. E. Hirsh, The application of statistical physics to evolutionary biology, Proceedings of the National Academy of Sciences 102 (2005) 9541–9546.

[89] D. M. McCandlish, A. Stoltzfus, Modeling evolution using the probability of fixation: history and implications, The Quarterly Review of Biology 89 (2014) 225–252.

[90] G. Sella, An exact steady state solution of Fisher’s geometric model and other models, Theoretical population biology 75 (2008) 30–4.

[91] D. D. Pollock, G. Thiltgen, R. A. Goldstein, Amino acid coevolution induces an evolutionary Stokes shift, Proceedings of the National Academy of Sciences of the United States of America 109 (2012) E1352–1359.

[92] G. Sella, D. H. Ardell, The impact of message mutation on the fitness of a genetic code, Journal of Molecular Evolution 54 (2002) 638–651.

[93] P. A. P. Moran, The survival of a mutant gene under selection. II, Journal of the Australian Mathematical Society 1 (1960) 485–491.

[94] P. R. Schimmel, D. Söll, Aminoacyl-tRNA synthetases: general features and recognition of transfer RNAs, Annual Review of Biochemistry 48 (1979) 601–648.

[95] R. W. Hamming, Error detecting and error correcting codes, The Bell System Technical Journal 29 (1950) 147–160.

[96] R. Rigler, U. Pachmann, R. Hirsch, H. G. Zachau, On the interaction of seryl-tRNA synthetase with tRNA Ser. A contribution to the problem of synthetase-tRNA recognition, European Journal of Biochemistry 65 (1976) 307–315.

[97] D. Riesner, A. Pingoud, D. Boehme, F. Peters, G. Maass, Distinct steps in the specific binding of tRNA to aminoacyl-tRNA synthetase. Temperature-jump studies on the serine-specific system from yeast and the tyrosine-specific system from Escherichia coli, European Journal of Biochemistry 68 (1976) 71–80.

[98] I. A. Vasil’eva, N. A. Moor, Interaction of aminoacyl-tRNA synthetases with tRNA: general principles and distinguishing characteristics of the high-molecular-weight substrate recognition, Biochemistry. Biokhimiia 72 (2007) 247–263.

[99] M. A. Savageau, R. R. Freter, On the evolution of accuracy and cost of proofreading tRNA aminoacylation, Proceedings of the National Academy of Sciences of the United States of America 76 (1979) 4507–4510.

[100] D. H. Ardell, G. Sella, No accident: genetic codes freeze in error-correcting patterns of the standard genetic code, Philosophical Transactions of the Royal Society of London. Series B, Biological Sciences 357 (2002) 1625–1642.

[101] G. Sella, D. H. Ardell, The coevolution of genes and genetic codes: Crick’s frozen accident revisited, Journal of Molecular Evolution 63 (2006) 297–313.

[102] A. Tietävöinen, On the Nonexistence of Perfect Codes over Finite Fields, SIAM Journal on Applied Mathematics 24 (1973) 88–96.

[103] F. I. Solov’eva, Perfect binary codes: bounds and properties, Discrete Mathematics 213 (2000) 283–290.

[104] H. Helgert, R. Stinaff, Minimum-distance bounds for binary linear codes, IEEE Transactions on Information Theory 19 (1973) 344–356.

[105] M. Best, A. Brouwer, F. MacWilliams, A. Odlyzko, N. Sloane, Bounds for binary codes of length less than 25, IEEE Transactions on Information Theory 24 (1978) 81–93.

[106] M. J. Rogers, D. Soll, Inaccuracy and the recognition of tRNA, Progress in Nucleic Acid Research and Molecular Biology 39 (1990) 185–208.

[107] F. Solov’eva, On perfect binary codes, Discrete Applied Mathematics 156 (2008) 1488–1498.

[108] W. Paulander, S. Maisnier-Patin, D. I. Andersson, Multiple mechanisms to ameliorate the fitness burden of mupirocin resistance in Salmonella typhimurium, Molecular Microbiology 64 (2007) 1038–1048.

[109] M. Ehrenberg, C. G. Kurland, Costs of accuracy determined by a maximal growth rate constraint, Quarterly Reviews of Biophysics 17 (1984) 45–82.

[110] J. Elf, D. Nilsson, T. Tenson, M. Ehrenberg, Selective Charging of tRNA Isoacceptors Explains Patterns of Codon Usage, Science 300 (2003) 1718–1722.

[111] J. Elf, M. Ehrenberg, Near-Critical Behavior of Aminoacyl-tRNA Pools in E. coli at Rate-Limiting Supply of Amino Acids, Biophysical Journal 88 (2005) 132–146.

[112] M. Johansson, M. Lovmar, M. Ehrenberg, Rate and accuracy of bacterial protein synthesis revisited, Current Opinion in Microbiology 11 (2008) 141–147.

[113] M. Kimura, Natural selection as the process of accumulating genetic information in adaptive evolution, Genetics Research 2 (1961) 127–140.

[114] J. Felsenstein, On the Biological Significance of the Cost of Gene Substitution, The American Naturalist 105 (1971) 1–11.

[115] J. Maynard Smith, The 1999 Crafoord Prize Lectures. The idea of information in biology, The Quarterly Review of Biology 74 (1999) 395–400.

[116] M. C. Donaldson-Matasci, C. T. Bergstrom, M. Lachmann, The fitness value of information, Oikos (Copenhagen, Denmark) 119 (2010) 219–230.

[117] J. J. Perona, Y.-M. Hou, Indirect Readout of tRNA for Aminoacylation, Biochemistry 46 (2007) 10419–10432.

[118] E. M. Novoa, M. Pavon-Eternod, T. Pan, L. Ribas de Pouplana, A Role for tRNA Modifications in Genome Structure and Codon Usage, Cell 149 (2012) 202–213.

[119] V. Bedian, Self-description and the origin of the genetic code, Bio Systems 60 (2001) 39–47.

[120] M. C. Cowperthwaite, L. A. Meyers, How Mutational Networks Shape Evolution: Lessons from RNA Models, Annual Review of Ecology, Evolution, and Systematics 38 (2007) 203–230.

[121] J. T.-F. Wong, Coevolution theory of the genetic code at age thirty, BioEssays: News and Reviews in Molecular, Cellular and Developmental Biology 27 (2005) 416–425.

[122] M. Di Giulio, An Autotrophic Origin for the Coded Amino Acids is Concordant with the Coevolution Theory of the Genetic Code, Journal of Molecular Evolution 83 (2016) 93–96.

[123] M. Lynch, K. Hagner, Evolutionary meandering of intermolecular interactions along the drift barrier, Proceedings of the National Academy of Sciences of the United States of America 112 (2015) E30–38.

[124] J. D. Watson, F. H. Crick, Genetical implications of the structure of deoxyribonucleic acid, Nature 171 (1953) 964–967.

[125] L. L. Cavalli-Sforza, Genes, peoples, and languages, Proceedings of the National Academy of Sciences of the United States of America 94 (1997) 7719–7724.

[126] A. Bouchard-Ct, D. Hall, T. L. Griffiths, D. Klein, Automated reconstruction of ancient languages using probabilistic models of sound change, Proceedings of the National Academy of Sciences of the United States of America 110 (2013) 4224–4229.

[127] S. Roman, Coding and Information Theory, Graduate Texts in Mathematics, Springer-Verlag, 1992.

[128] M. Di Giulio, Was it an ancient gene codifying for a hairpin RNA that, by means of direct duplication, gave rise to the primitive tRNA molecule?, Journal of Theoretical Biology 177 (1995) 95–101.

[129] J. Widmann, M. Di Giulio, M. Yarus, R. Knight, tRNA creation by hairpin duplication, Journal of Molecular Evolution 61 (2005) 524–530.

[130] S. Karlin, S. F. Altschul, Methods for assessing the statistical significance of molecular sequence features by using general scoring schemes, Proceedings of the National Academy of Sciences of the United States of America 87 (1990) 2264–2268.

[131] D. T. Gillespie, Exact stochastic simulation of coupled chemical reactions, The Journal of Physical Chemistry 81 (1977) 2340–2361.

[132] P. Shah, Y. Ding, M. Niemczyk, G. Kudla, J. B. Plotkin, Rate-Limiting Steps in Yeast Protein Translation, Cell 153 (2013) 1589–1601.

